# Optimizing genomic sampling for demographic and epidemiological inference with Markov decision processes

**DOI:** 10.1101/2025.06.30.662264

**Authors:** David A. Rasmussen, Madeline G. Bursell, Frank Burkhart

**Affiliations:** Dept. of Entomology and Plant Pathology, North Carolina State University, Raleigh, NC, USA; Bioinformatics Research Center, North Carolina State University, Raleigh, NC, USA

## Abstract

Inferences from population genomic data provide valuable insights into the demographic history of a population. Likewise, in genomic epidemiology, pathogen genomic data provide key insights into epidemic dynamics and potential sources of transmission. Yet predicting what information will be gained from genomic data about variables of interest and how different sampling strategies will impact the quality of downstream inferences remains challenging. As a result, population genomics largely lacks theory to guide decisions on how best to sample individuals for genomic sequencing. By adopting a sequential decision making framework, we show how Markov decision processes (MDPs) can be applied to jointly model a population’s dynamics along with the sampling process. Critically, these MDPs allow us to compute the expected long-term value of sampling in terms of information gained about estimated variables. This in turn allows us to very efficiently explore and identify optimal sampling strategies. To illustrate our framework, we develop MDPs for three common demographic and epidemiological inference problems: estimating population growth rates, minimizing the transmission distance between sampled individuals and estimating migration rates between subpopulations. In each case, the MDP allows us to identify optimal sampling strategies that maximize the information gained from genomic data while minimizing costs.

## Introduction

The ability to sequence many individual genomes from a population has greatly expanded our ability learn about a population’s demographic history, including changes in population size (1; 2), population structure (3; 4), and ancestry (5; 6; 7). Genomic epidemiology provides a remarkable example of this trend, with pathogen genomes now being routinely analyzed to infer changes in epidemic dynamics (8; 9) and reveal potential pathways of transmission (10; 11; 12). Despite these advances, it remains generally unclear how best to sample individuals for the purposes of genomic sequencing and downstream demographic or epidemiological inference (13; 14). Because it is usually only feasible to sample a small fraction of the total population, questions about sampling design inevitably arise, including: who to sample, what sample sizes are sufficient, and how representative must sampling be to provide reliable inferences?

Unfortunately, unlike other areas of modern statistical inference (15; 16; 17), population genomics and genomic epidemiology largely lack theory to guide decisions on how best to sample and sequence individual genomes (herein we refer to both sampling and sequencing simply as *sampling*). In absence of such guidance, sampling is often performed based on simple heuristics or opportunistically based on available resources (18; 19). As a result, sampling can be highly biased and non-representative. These sampling biases can in turn bias downstream demographic inferences, which may be quite sensitive to violations of assumptions about how individuals are sampled (20; 21; 22; 23; 24).

At the same time, optimizing sampling strategies in population genomics can be challenging for several reasons (Fig 1). First, the optimal sampling strategy may be highly dependent on a population’s dynamics, which may not be fully observable (Fig 1A). Second, the information gained from sampling impacts may be highly dependent on past sampling effort. Optimal strategies therefore need to reflect what data is already available and the current level of uncertainty surrounding estimated variables (Fig 1B). Third, what information will be gained from new data is often highly unpredictable. Sampling more may in some cases improve estimates, but in other cases may only exacerbate existing biases (Fig 1C). As we elaborate on below, these challenges largely stem from a more fundamental problem: the information gained from genomic data almost always depends on how sampled individuals are related to one another through their shared ancestry or phylogenetic history, yet these relationships cannot be known *a priori* before sampling. Ultimately, this inability to predict the informational value of genomic data makes it difficult to answer even the most pragmatic questions about sampling, such as what is an appropriate sample size?

**Fig 1.**
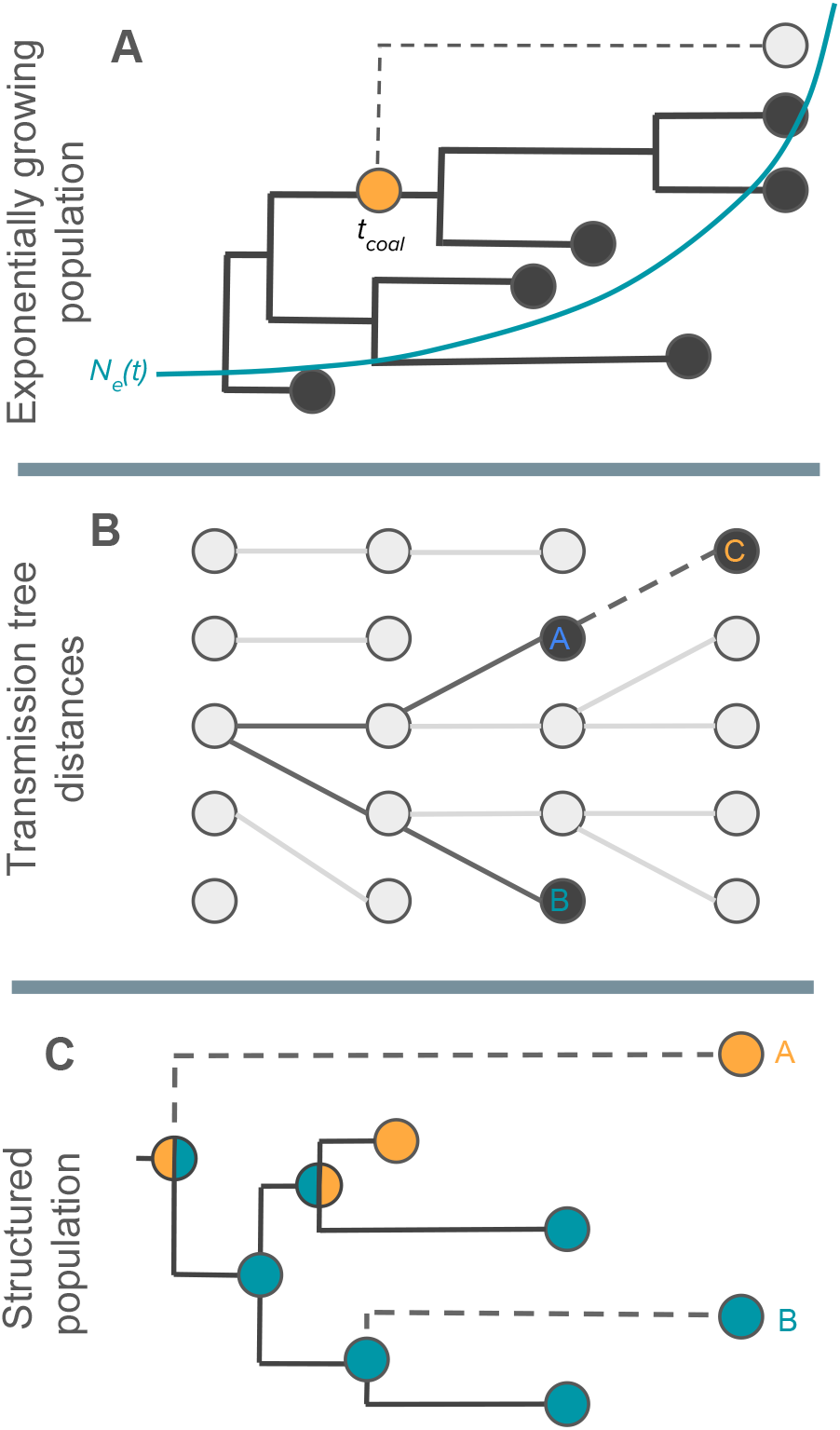
Challenges associated with optimizing genomic sampling. The three examples illustrate how the informational value of sampling can depend on the ancestral relationships among sampled individuals. **A:** In coalescent-based inference tasks such as inferring past changes in population size, the ancestral relationships among sampled individuals will determine when coalescent events are observed, which will in turn depend on past population sizes *N*_*e*_(*t*). For example, sampling at present may provide little information about recent population sizes if all sampled lineages coalesce deep in the past. **B:** In genomic epidemiology, interest often lies in reconstructing transmission trees to learn about potential transmission pathways. Sampling individuals linked by one or a small number of intervening transmission events (i.e. short transmission distances) can provide more information about who may have infected whom. However, the distance between sampled individuals may depend on past sampling. Here, the transmission distance between individual C and its nearest sampled neighbor depends entirely on who was previously sampled. **C**: In phylogeography, the movement of individuals is reconstructed based on the probable location of ancestral nodes. However, different patterns of movement can be inferred depending on exactly which individuals are sampled from each population. In this case, sampling individual A reveals an additional migration event that could dramatically alter the most probable root state, and therefore the presumptive source population, whereas sampling individual B provides little additional information about movement.

**Fig 1.**
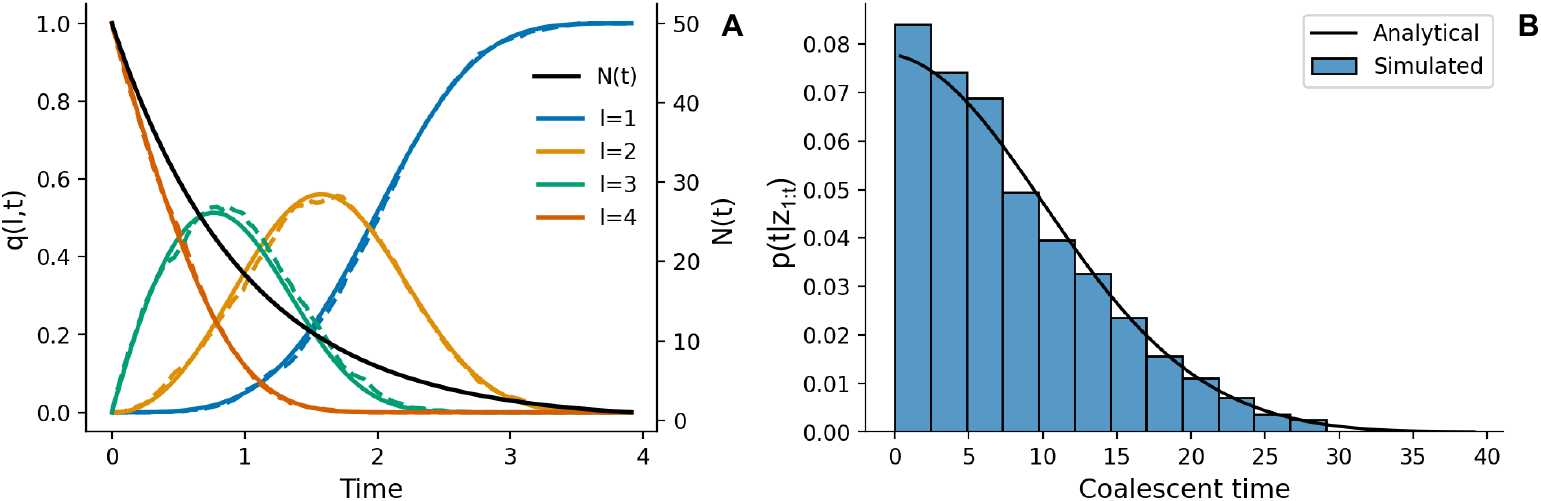
Computing the LTT and coalescent time density for the exponential growth MDP. **A:** The probability density *q*(*l, τ*) for the number of lineage *l* through time (solid lines) compared with Monte Carlo simulations of the coalescent process (dashed lines) with 4 lineages sampled at present. **B**: The probability density *p*(*τ* |*z*_1:*t*_) for the time *t*_*coal*_ at which a sampled lineage coalesces with the rest of the sample compared against Monte Carlo simulations. Here, *p*(*τ* | *z*_1:*t*_) was computed using the LTT density *q*(*l, τ*) shown in B.

As a step towards optimizing genomic sampling, we pose the problem of sampling design within a sequential decision making framework (25; 16). This framework allows us to jointly model the sampling process together with the underlying ecological or epidemic dynamics of a population as a Markov decision process (MDP). MDPs are widely used to optimize the control of dynamical systems (26; 27; 28), and are increasingly being applied to problems such as optimizing epidemic control and antibiotic drug treatment strategies (29; 30; 31). While our interest lies not in directly controlling the dynamics of a population, we show how the well-developed theoretical machinery of MDPs can be adapted to efficiently predict the long-term reward or value of a given sampling strategy in terms of the information gained about an estimated parameter or variable. MDPs can therefore guide and structure the search towards strategies that optimize the expected value of sampling.

While our framework has much in common with standard MDPs, extending them to genomic sampling presents several difficulties. We therefore begin with a general review of MDPs for sequential decision making problems before explaining how MDPs can be adapted to optimize genomic sampling. We then apply our framework to three common scenarios in population genomics: estimating growth rates of a growing population, minimizing transmission distances between sampled individuals and estimating migration rates between subpopulations. In the Results, we explore optimal sampling strategies for each scenario and compare the learned strategies against brute force searches to demonstrate that the strategies identified by MDPs are in fact globally optimal.

## Models and methods

### Markov decision processes

We first describe the core components of a generic Markov decision process (MDP). In a MDP, an agent representing a decision maker interacts with its environment through a series of actions. At each decision time point, the agent receives a reward based on the chosen action and the state of the environment, which may depend on the agent’s previous actions. In most applications, the goal is for the agent to learn how to make actions that maximize the total or long-term reward.

Adopting standard notation (27; 32), let *s*_*t*_ ∈ 𝒮 and *a*_*t*_ ∈ 𝒜 represent the state of the environment and the action the agents chooses at time *t*, respectively. The dynamics of the environment are described by a set of transition probabilities *p*(*s*_*t*+1_|*s*_*t*_, *a*_*t*_), which provide the probability of the state changing from state *s*_*t*_ at time *t* to *s*_*t*+1_ at time *t* + 1 given that action *a*_*t*_ was taken by the agent. The actions the agent takes are determined by a policy function *π*(*a*|*s*), which may be either deterministic or stochastic. For any action *a* taken from any state *s*, the agent receives a reward *r* ∈ ℛ with probability *p*(*r*|*s, a*). The expected reward the agent receives for any state-action pair is given by the reward function:

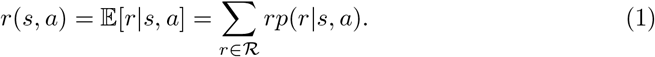

We assume throughout that decisions are made at a fixed sequence of time points *t* = 1, …, *N* over a finite horizon ending at *t* = *N*. However the underlying dynamics of the Markov process may evolve on a different timescale in discrete or continuous time depending on the application. To distinguish between these timescales, we let *τ*_*t*_ represent the actual time at which decision *t* is made. The time elapsed between two successive decision points Δ*τ* = *τ*_*t*+1_ − *τ*_*t*_ is arbitrary and not necessarily constant.

We are generally interested in finding an optimal policy function *π*^*∗*^ that maximizes the long-term rewards the agent receives at all future times. Here, the optimal policy *π*^*∗*^ is formally defined as the strategy that maximizes the expected cumulative sum of future rewards:.

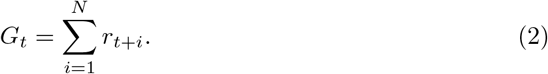

In order to learn the optimal policy, we need to be able to compute the expected value or long-term reward of a given policy. The expected value of a given policy *π* starting in state *s*_*t*_ can be computed recursively using the Bellman equations (25; 26):

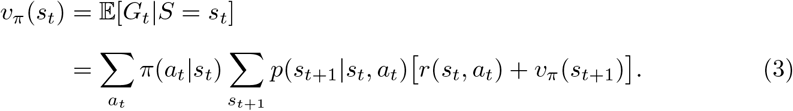

We can also write the Bellman equations in terms the expected value of taking action *a*_*t*_ in state *s*_*t*_ under a given policy *π*:

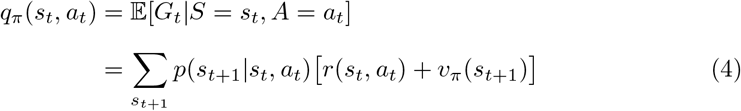

The value function *q*_*π*_(*s*_*t*_, *a*) enables us to compute the total expected reward that will be gained over time if a given action is taken at the current time and we continue to act according to the policy *π*. Being able to compute these expected long-term values is vitally important for identifying actions that are globally optimal and not just locally optimal in terms of maximizing the immediate rewards received upon taking an action. As we shall see, this becomes especially important for learning optimal genomic sampling strategies where the long-term rewards of sampling at present will often depend on past and future sampling decisions.

### Learning optimal strategies using dynamic programming

The value functions presented in (3) and (4) can be efficiently evaluated using dynamic programming (33). To evaluate the expected value of a given policy *π*, we use an iterative policy evaluation algorithm (32). Starting from an initially arbitrary estimate of the value *V* (*s*_*t*_) for each state *s*_*t*_ ∈ 𝒮, each iteration of the algorithm updates the estimate of *V* (*s*_*t*_) using the expected value of successor states *V* (*s*_*t*+1_) obtained from previous iterations. This procedure is iterated until the algorithm converges on a stable estimate of *V*.

To identify the optimal policy, we use a backward induction or value iteration algorithm (27; 32). This algorithm combines iteratively updating expected values (as in policy evaluation) with improvements to the policy. While the policy is not explicitly updated, the values are always updated with the expected value of the action that would maximize long-term expected value, such that the policy implicitly improves over time. This procedure is iterated until the algorithm converges on a stable estimate of *V* where the policy can no longer be improved. Given the optimized value function *V*, we can then find the optimal action *a*^*∗*^ to take from any state *s*_*t*_: *a*^*∗*^ = arg max_*a*_[*q*_*π*_(*s*_*t*_, *a*)]. Identifying *a*^*∗*^ for all possible states therefore provides the optimal policy. Pseudocode for the policy evaluation and value iteration algorithms is provided in S1 Appendix.

### Markov decision processes for optimal genomic sampling

Before considering specific MDPs for genomic sampling, we consider the general motivation behind our approach. Overall, our goal is to maximize the amount of information we gain from genomic data about estimated variables of interest, while also minimizing the total cost of sampling. How much information we gain from sampling will generally depend on how sampled individuals are related to one another through their shared ancestry of phylogenetic history. For example, how much information is gained from sampling an individual under standard coalescent models will depend on when coalescent events occur in the ancestry of the sample (Fig 1A). While it may be possible to reconstruct these phylogenetic relationships from genomic data, we do not know how sampled individuals will be related to one another prior to sampling, and thus cannot directly predict what information will be gained from sampling. This is indeed one of the major challenges of designing optimal sampling strategies: we generally don’t know the value of sampling until we actually sample.

Our MDP framework attempts to circumvent this problem by finding the expected value of sampling without *a priori* knowledge of how sampled individuals will be related. In essence, we compute expected values by marginalizing or integrating over all possible ancestral or phylogenetic histories of the sample. When computing the expected value of a newly sampled individual, we consider where and when this individual might “attach” or coalesce with the rest of the sample. We can then compute the expected value of the new sample by integrating over all possible attachment points, weighting each by what information would be gained from observing an event at the time and/or location.

This MDP framework can be thought of as a special case of the more general MDPs considered above. The Markov process describes how the underlying population dynamics gives rise to the phylogeny relating the sampled individuals. On top of this Markov process is a sequential decision making process that determines who or how many individuals are sampled at each decision time point. In this case:

- The state represents the *sample configuration*, typically the number or set of individuals sampled at each time.
- Actions update the state through sampling events.
- The action policy represents the sampling strategy.
- The rewards reflect information gained from the sample about estimated quantities of interest.

In order to find the optimal sampling strategy, we need to be able to evaluate the long-term expected value of a given sampling or action policy. This in turn requires us to compute the expected reward *r*(*s, a*) of taking a particular action from a given state. In the relatively simple MDPs we consider next, these expected rewards can be computed analytically. While it may not always be possible to compute expected rewards analytically, other strategies such as Monte Carlo simulations can be performed to approximate the rewards. Either way, once we can compute the the expected rewards, we can combine the value functions in (3) and (4) with dynamic programming to learn the optimal sampling policy that maximizes long-term expected rewards.

### Exponential growth coalescent MDP

We first consider a MDP for sampling individuals under the coalescent with exponential growth (1; 34). Fig 2 provides a schematic representation of the MDP. Our overall goal is to optimize sampling in order to maximize information about the estimated population growth rate *γ* while minimizing the total cost of sampling.

**Fig 2.**
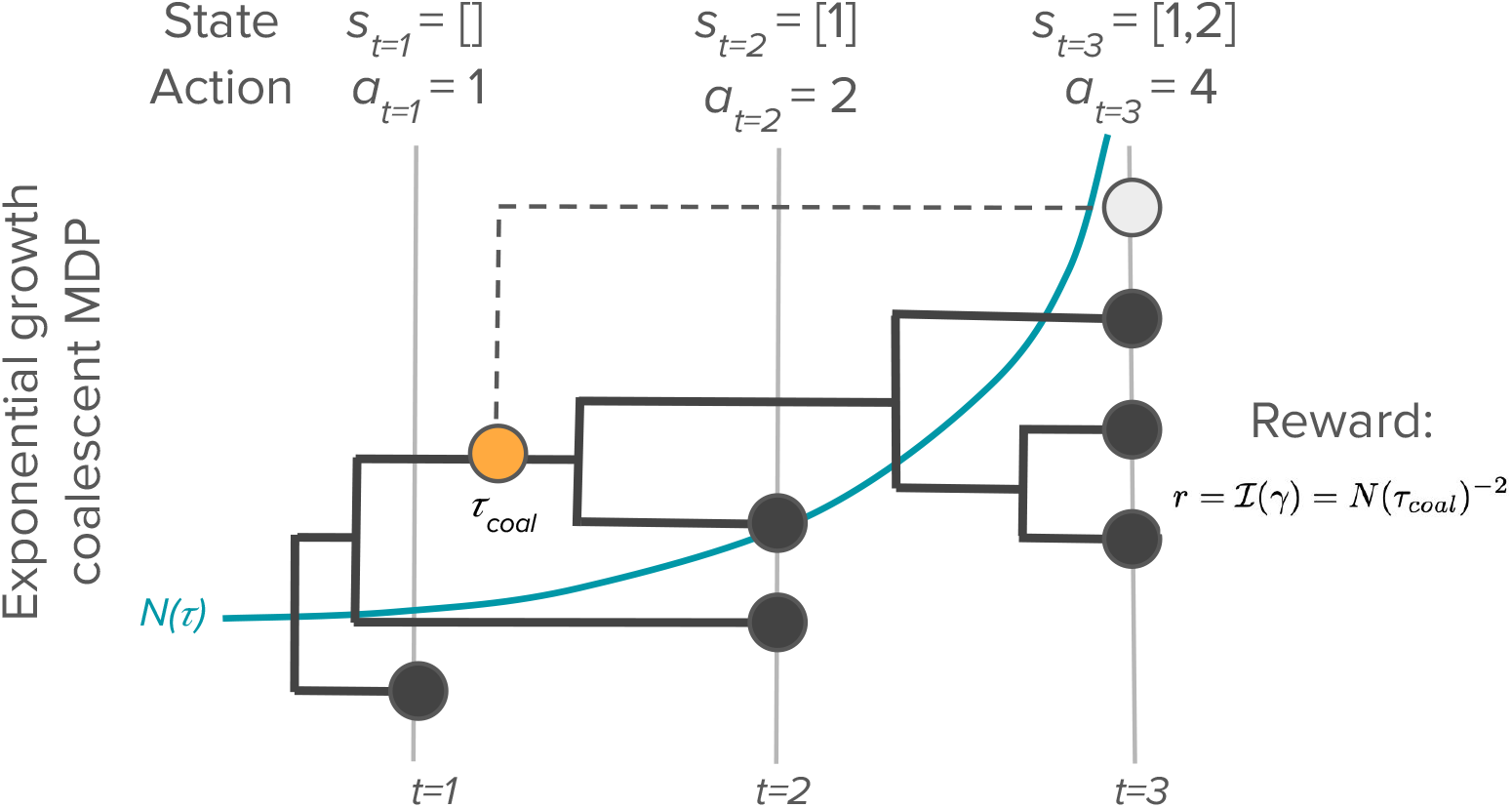
The exponential growth coalescent MDP. At each decision time *t*, the agent decides what action *a*_*t*_ to take in terms of how many individuals to sample. The state *s*_*t*_ is determined by the number of samples taken at each previous time point up to *t*. The reward function quantifies how much Fisher information about the growth rate *γ* is gained from a sampling action. Here, we illustrate computing the reward for the light gray node sampled at *t* = 3. The amount of Fisher information gained from sampling this individual will depend on the population size *N* (*τ*_*coal*_) at the time *τ*_*coal*_ when the sampled lineage coalesces with the rest of the sample. Because we do not know when *τ*_*coal*_ will occur before sampling, we compute the expected reward by integrating over all possible coalescent times.

**Fig 2.**
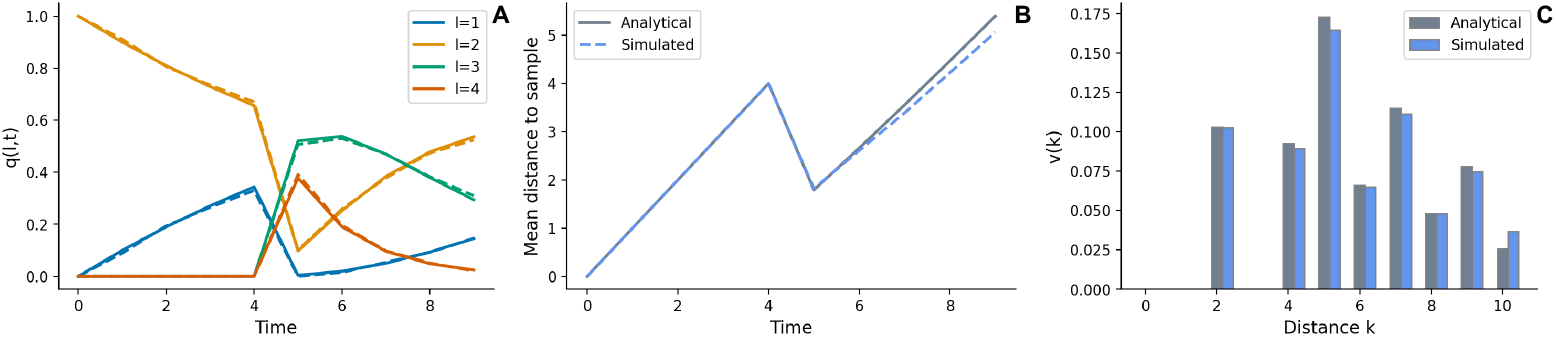
Probability densities used to compute expected rewards under the transmission tree MDP. **A**: The probability density *q*(*l, t*) for the number of lineages through time (solid lines) compared with Monte Carlo simulations of transmission trees (dashed lines). **B**: The mean ancestor-to-sample distance computed based on the probability density *m*(*k, t*) (solid line) compared with Monte Carlo simulations (dashed line). **C**: The probability density *v*(*k* |𝒵_1:*t*_) that a sample has its nearest sampled neighbor at transmission distance *k*. In all cases, *N* (0) = 10 with with 2 lineages sampled at present (*t* = 0) and 2 lineages sampled at *t* = 4.

#### Markov process

We assume that the population grows deterministically in continuous time such that the population size *N* (*τ*) at time *τ* in the past is given by:

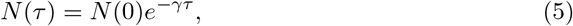

where *N* (0) is the population size at present.

While the population dynamics are deterministic, the coalescent model tracks the ancestry of sampled individuals backwards through time as a stochastic process.

Lineages ancestral to sample coalesce at a rate *λ*(*τ*) determined by the time-varying population size *N* (*τ*) and the number of lineages ancestral to the sample *k*:

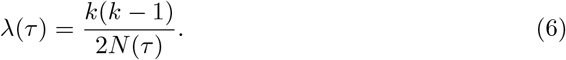

The probability of a coalescent event occurring at time *τ* in the past follows an exponential probability density:

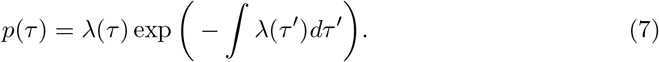

#### Action space

On top of the underlying coalescent process, there is a sequential decision making process where at each time point *t* the agent decides how many individuals *a*_*t*_ = *z*_*t*_ to sample according to the policy *π*(*a*|*s*).

Note that while time in the coalescent model goes backwards from *τ* = 0 at present, we order the decision time points in forward time from *t* = 1 at the earliest decision time (furthest in the past) to the last decision time point at *t* = *N*. We use *τ*_*t*_ to refer to the actual time at which decision time point *t* in units of backwards coalescent time.

#### State space

The state *s*_*t*_ of the system is determined by the sample configuration *z*_1:*t*_ = *z*_1_, *z*_2_,…, *z*_*t*_, the number of individuals sampled at each decision time up to time *t*. The population size *N* (*τ*) is assumed known from the deterministic population dynamics.

#### Reward function

Sampling each individual adds one additional coalescent event to the phylogenetic tree. To quantify the information gained about *γ* from each coalescent event, we use Fisher information (35; 36). With constant population size *N*, the waiting times between coalescent events are independently and exponentially distributed with rate parameter *λ* ∝ 1*/N*. The Fisher information ℐ(*N*) about *N* provided by each coalescent event is therefore the same as the observation of an exponentially distributed random variable ℐ(*N*) = *λ*^2^ = (1*/N*)^2^ = *N* ^*−*2^ (37). Because the waiting time between events are independent, the information provided by each sample is additive such that the total Fisher information provided by *m* samples is ℐ(*N*) = *mN* ^*−*2^.

For an exponentially growing population, the waiting times between coalescent events remain independently distributed random variables, but they are no longer exponentially distributed because the coalescent rate increases as *N* decreases into the past. However, we assume that ℐ(*γ*) is still proportional to the population size at the time *τ* at which coalescent events occurs such that ℐ(*γ*) = *N* (*τ*)^*−*2^. Simulation results suggest this is an excellent approximation (see Fig 6C). Thus, coalescent events occurring further in the past when the population is small will carry more information about the growth rate than events closer to present.

#### Expected rewards

The value of sampling each individual depends on when the sampled lineage coalesces with the other lineages in the sample, which we cannot observe before sampling. Therefore, in order to compute the expected value of sampling a given individual, we track the probability density *p*(*τ* |*z*_1:*t*_) for the time *τ* at which the sampled individual coalesces with the other sampled lineages, conditional on the current sample configuration *z*_1:*t*_. Full details on how *p*(*τ* |*z*_1:*t*_) is computed are provided in S2 Appendix.

Given *p*(*τ* |*z*_1:*t*_), the expected value of sampling one individual in terms of the Fisher information ℐ(*γ*) can then be computed by integrating over the unknown time at which the newly sampled lineage coalesces with the other sampled lineages:

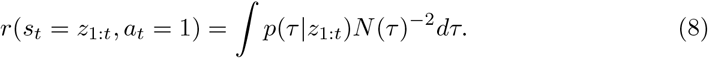

Because the reward of sampling an individual at time *t* will depend on how many other individuals are sampled at time *t*, we rewrite *z*_1:*t*_ as the tuple (*z*_1:*t−*1_, *z*_*t*_) to distinguish between samples taken in previous generations and the present time *z*_*t*_. The full expected value of sampling *z*_*t*_ individuals conditional on *z*_1:*t−*1_ can then be computed iteratively:

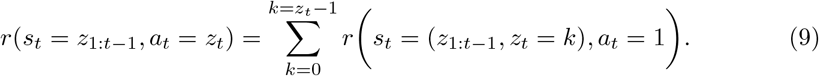

### Transmission tree distance MDP

Minimizing the transmission distance, or the number of intervening transmission events between sampled individuals, is often a goal in genomic epidemiology when interest lies in *linking* infected individuals based on genetic data as a means to infer who might have infected whom (13; 38; 39). The goal of our second MDP therefore is to minimize the transmission distance between sampled individuals in a transmission tree (Fig 3). Nodes in the transmission tree represent infected individuals and edges linking nodes describe who infected whom through transmission events. We define the transmission distance *d*_*ij*_ to be the number of edges traversed along the shortest path connecting nodes *i* and *j* in a transmission tree 𝒯. A direct transmission pair will therefore have *d*_*ij*_ = 1.

**Fig 3.**
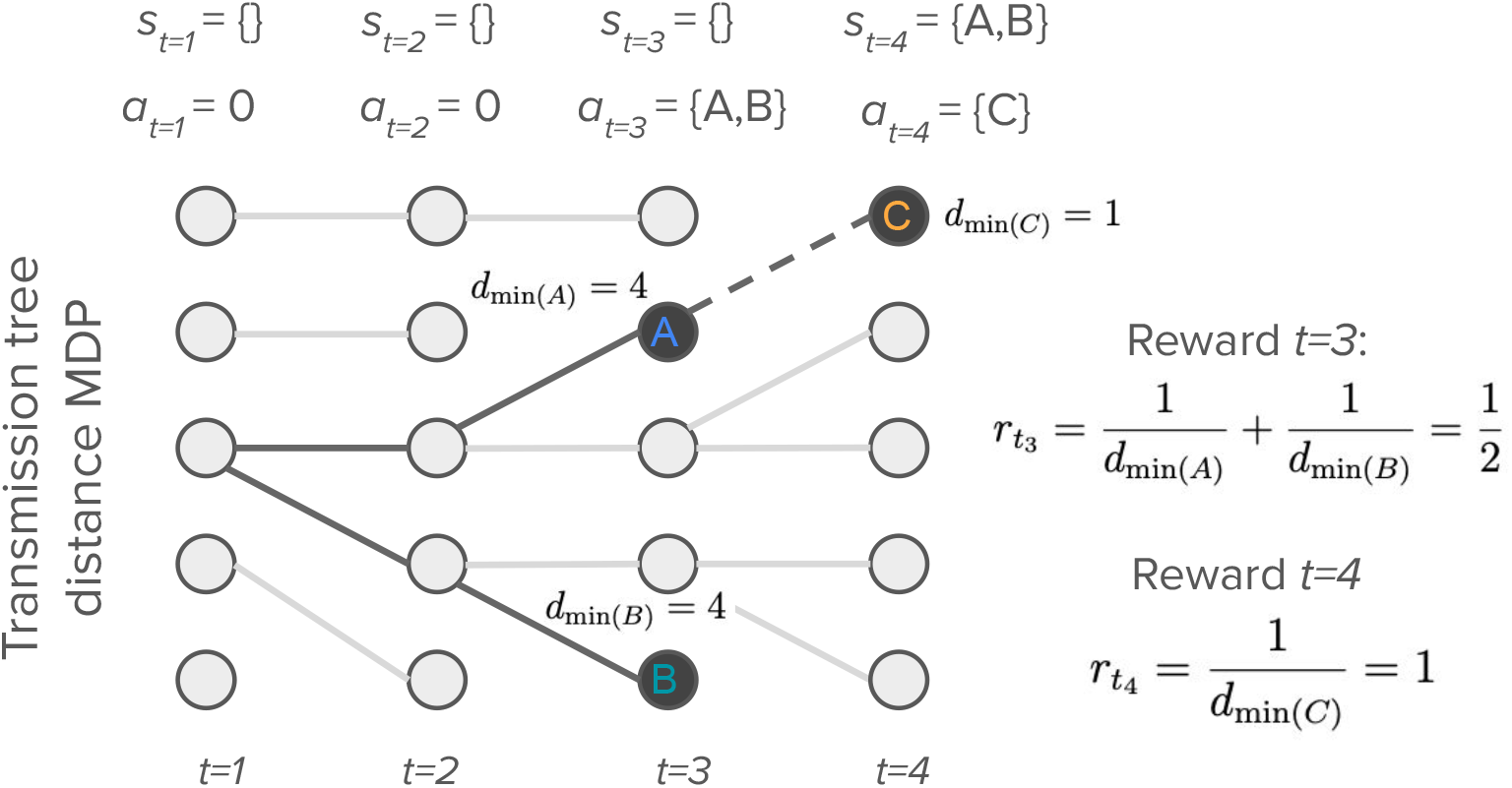
The transmission tree distance MDP. At each decision time *t*, the agent decides what action *a*_*t*_ to take in terms which individuals to sample. The state *s*_*t*_ is determined by the set of all individuals sampled up to *t*. The reward function quantifies the reward gained from sampling individuals at a given transmission distance *d*_*min*(*i*)_. Here, we illustrate computing the rewards based on inverse distances, in which the reward received from sampling an individual is equal to the inverse of the transmission distance between the newly sampled node and its nearest sampled neighbor.

**Fig 3.**
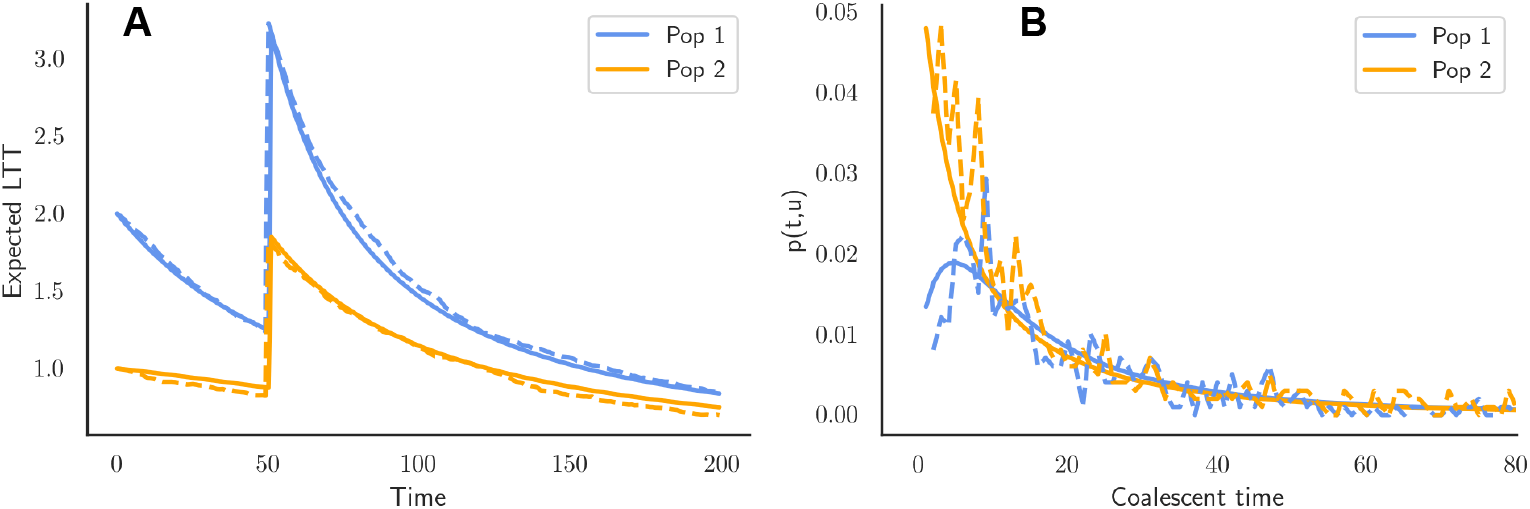
The LTT and coalescent densities for the structured coalescent MDP. **A**: The expected number of lineages in each population through time computed from the LTT density *q*(*k, l, τ*) (solid lines) compared with Monte Carlo simulations of the coalescent process (dashed lines). In this case, two lineages are sampled from population 1 and one lineage is sampled from population 2, both at present and at time *τ* = 50 in the past. **B**: The probability density *p*(*τ, u* | *z*_1:*t*_) for the time *t*_*coal*_ at which a lineage coalesces with a parent lineage in population *u* = 1 or population *u* = 2. The coalescent density computed using Eq 19 (solid lines) is compared against Monte Carlo simulations of the coalescent process (dashed lines). Here, two lineage are assumed to have already been sampled in each population at present (*t* = 0), and *p*(*τ, u* | *z*_1:*t*_) is given for a fifth lineage sampled in population 2.

#### Markov process

A stochastic branching process generates topology of the transmission tree. For simplicity, we assume infections occur in non-overlapping generations with *I*(*t*) infections present in generation *t*. At each generation, the parent node of each newly infected node is chosen by randomly sampling nodes in the previous generation with uniform probability and with replacement. The epidemic dynamics of how *I*(*t*) changes through time is assumed to be deterministic.

#### Action policy

At each generation *t*, the agent decides how many individuals *z*_*t*_ to sample according to the policy *π*(*a*|*s*) and then randomly samples a set *Ƶ*_*t*_ of individuals. Because the transmission distance between samples will depend on exactly which individuals are sampled, we assume that the agent remembers the full set of individuals *Ƶ*_*t*_ sampled at generation *t*.

#### State space

The state *s*_*t*_ of the system is given by the sample configuration *Ƶ*_1:*t*_ = *Ƶ*_1_, *Ƶ*_2_,…, *Ƶ*_*t*_ composed all individuals sampled up to generation *t*. Note that in this case we assume decision time points occur on the same timescale as generations in the transmission process.

#### Reward function

The reward the agent receives for sampling is determined by the transmission distance between each sampled individual and its nearest sampled neighbor in the transmission tree. We denote the transmission distance between *i* and its nearest sampled neighbor as *d*_min(*i*)_. For a given set of sampled individuals *Ƶ*:

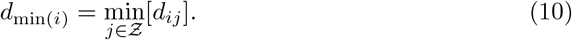

We explore two reward functions that differ in how the rewards depend on *d*_min(*i*)_. We refer to the first as the *inverse distance* reward function because it assumes the value of a sample is inversely proportional to its distance to its nearest sampled neighbor:

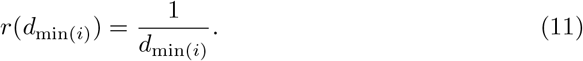

We refer to the second as the *direct pairs* reward function as it assumes that only direct transmission pairs are of interest. In this case:

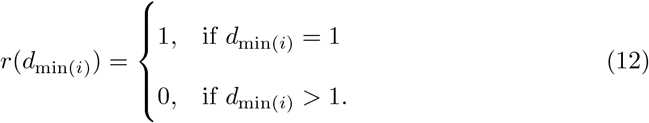

#### Expected rewards

In order to compute the expected value of sampling a given individual, we first compute the probability *v*(*k*|*Ƶ*_1:*t*_) that the transmission distance between the individual and its nearest sampled neighbor in *Ƶ*_1:*t*_ is *k*. In brief, computing *v*(*k*|*Ƶ*_1:*t*_) requires integrating over all possible transmission trees, all possible locations (i.e. nodes) at which the newly added sample could attach in these trees, and the corresponding distances between these possible attachment nodes and the sample. Full details on how *v*(*k*|*Ƶ*_1:*t*_) is computed are provided in S3 Appendix.

To compute the expected value of a sample, we sum the rewards received at each possible transmission distance *k* weighted by the probability *v*(*k*|*Ƶ*_1:*t*_) that the sample has a nearest sample at that distance. Assuming we sample a single individual *i*, the expected reward is:

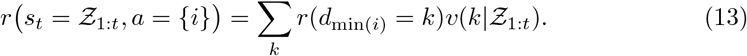

However, an additional complication arises here because sampling at present may change the distance of previous samples to their nearest sample. For example, we may sample a direct child of an individual sampled one generation in the past, converting their transmission distance to one. We therefore need to compute the total expected value of sampling all previously and newly sampled individuals *Ƶ*_1:*t*_ = *Ƶ*_1:*t−*1_ ∪ *Ƶ*_*t*_:

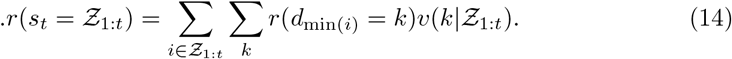

To compute the total expected value gained from sampling a set of individuals *Ƶ*_*t*_ at present, we then compare the rewards expected with and without the new samples in *Ƶ*_*t*_:

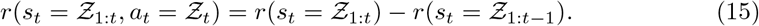

### Structured coalescent MDP

Finally, we consider a MDP for sampling individuals under a structured coalescent model (Fig 4) where individuals can migrate between two populations (3; 40). For an infectious pathogen, migration events may correspond to transmission events between different host populations. Our goal is to optimize sampling in order to maximize information about the estimated migration rates.

**Fig 4.**
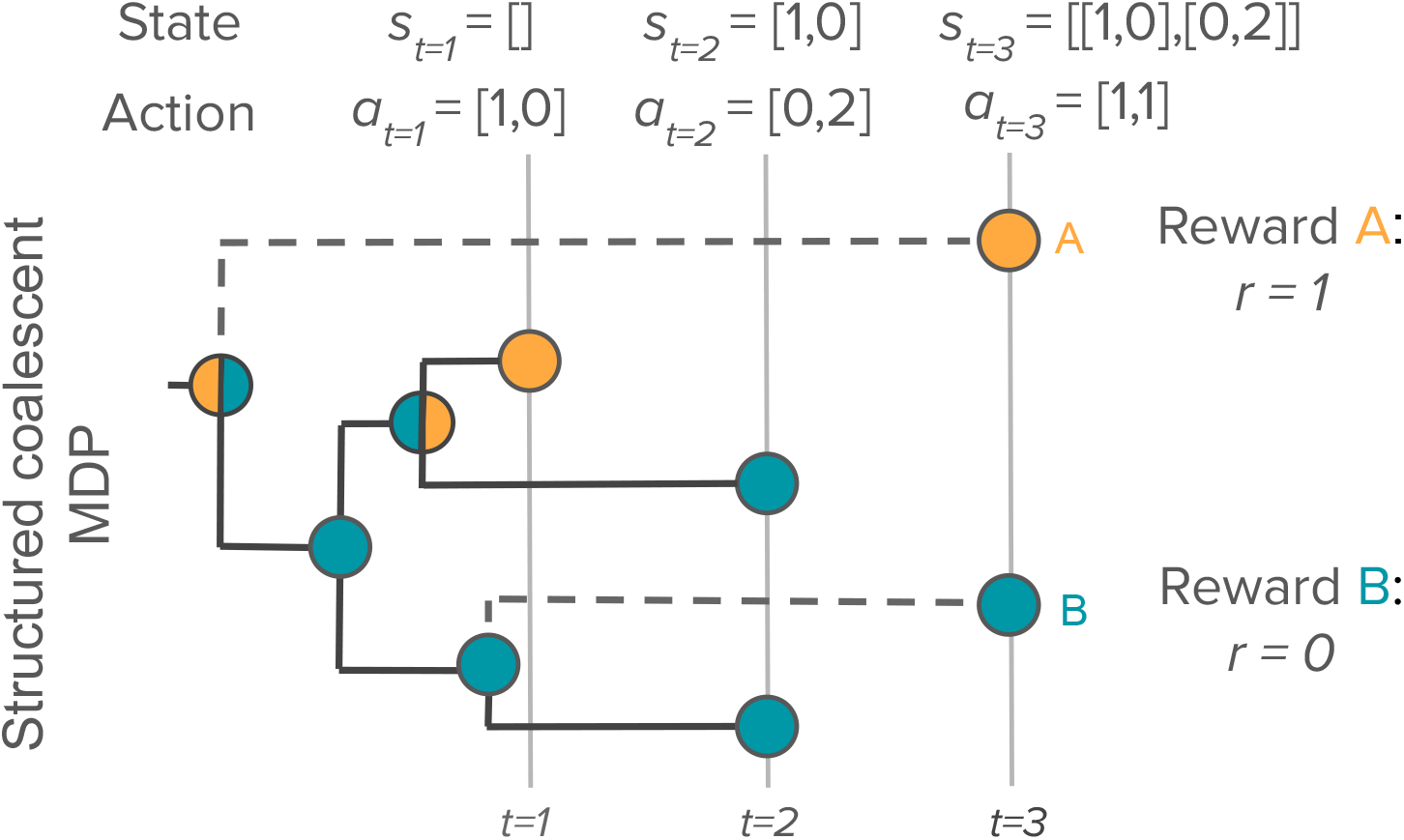
The structured coalescent MDP. At each decision time *t*, the agent decides on an action *a*_*t*_ that determines how many individuals are sampled from each population. The state *s*_*t*_ is determined by the number of samples taken from each population at all time points up to *t*. The reward function quantifies how many coalescent events occur between populations as a proxy for the information gained about the estimated migration rates. Because we are interested in estimating migration rates between populations, coalescent events occurring between as opposed to within populations will provide information about migration rates. Thus sampling individual *A* provides a reward *r* = 1 as it adds a coalescent event occurring between populations, whereas sampling individual *B* does not and therefore provides a reward *r* = 0. Because we do not know when and where sampled lineages will coalesce, the expected rewards are computed by integrating over all possible coalescent times and possible states of the parent and child lineages.

#### Markov process

The coalescent process underlying the MDP is similar to the exponential growth coalescent model except now individuals can migrate between populations. We assume that migration events occur through birth or transmission events between populations and that migrations cannot occur independently of births. The migration rate *m*_*i,j*_ is defined as the probability that a child in population *j* has a parent in population *i* in the previous generation (41). The proportion of individuals descended from parents in the same population is therefore 1 − *m*_*i,j*_. The size of each population *N*_*i*_(*τ*) may deterministically vary through time independent of the other population.

Going backwards through time, pairs of lineages coalesce at a rate *λ*_*ij*_(*τ*) that depends on what populations *i* and *j* they reside in:

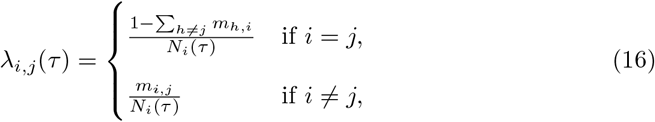

where here *i* denotes the state of the parent lineage and *j* the state of the child lineage.

#### Action space

At each decision time point *t*, the agent decides how many individuals *a*_*t*_ = [*z*_*i,t*_, *z*_*j,t*_] to sample in populations *i* and *j* according to the policy *π*(*a*|*s*). To simplify the action space, we again consider a small set of sampling fractions in each population. The set of all possible actions 𝒜(*t*) therefore includes all pairwise combinations of sampling fractions between populations.

#### State space

The state *s*_*t*_ of the system is determined by the sample configuration *z*_1:*t*_, where now *z*_*t*_ = [*z*_*i,t*_, *z*_*j,t*_].

#### Reward function

In structured coalescent models, inferences about migration rates are drawn based on information about the probable state (e.g. population or location) of lineages at the time of coalescent events (4; 42; 43). For example, if two coalescing lineages have a high probability of being in different populations, we can infer that the coalescent event likely occurred at a birth or transmission event between populations as opposed to an event within either population. Combining information about the probable ancestral state of lineages at coalescent events across a phylogeny allows for migration rates to be estimated while allowing for uncertainty in the ancestral states.

Based on the preceding logic, we reason that the information gained from a phylogeny about migration rates will depend on the number of coalescent events that occur between as opposed to within populations. For example, if we are interested in estimating the migration rate *m*_*i,j*_, we expect that the number of coalescent events *C*_*i,j*_ that occur where the parent lineage is in population *i* and one of the child lineages is in population *j* will be a reasonable proxy for the information gained about *m*_*i,j*_.

For our two-population model, we estimate the migration rates *m*_1,2_ and *m*_2,1_ corresponding to migration in both directions. Our reward function therefore quantifies the number of migration events that occur in either direction:

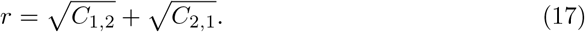

We square root transform the number of events as this was found to better predict the error in estimated migration rates in practice (see Fig 8).

#### Expected rewards

As before, sampling each individual adds one additional coalescent event to the phylogeny. However, we do not know when this coalescent event will occur nor the states of the lineages involved at the time of the event. In order to compute the expected value of sampling a given individual, we therefore track the probability density *p*(*τ, u, v*|*z*_1:*t*_) for the time *τ* of the coalescent event along with the state of the parent lineage *u* and the state of the sampled child lineage *v* at the coalescent event, conditional upon the current sample configuration *z*_1:*t*_. Full details on how *p*(*t, u, v*|*z*_1:*t*_) is computed are provided in S4 Appendix.

Given *p*(*τ, u, v*|*z*_1:*t*_), the expected value of sampling one individual in population *I* (*a*_*i,t*_ = 1) can by computed by considering the total probability that the coalescent event added by sampling occurs between lineages in different populations (*u* = 1 and *v* = 2 or *u* = 2 and *v* = 1) while integrating out the unknown coalescent time:

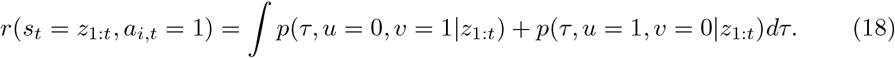

When sampling more than a single individual at one time, we rewrite *z*_1:*t*_ as the tuple (*z*_1:*t−*1_, *z*_*i,t*_, *z*_*j,t*_) to distinguish between samples taken in previous generations and the samples taken at present in different populations. The full expected value of sampling *z*_*t*_ = (*z*_*i,t*_, *z*_*j,t*_) individuals in each population conditional on *z*_1:*t−*1_ can then be computed iteratively:

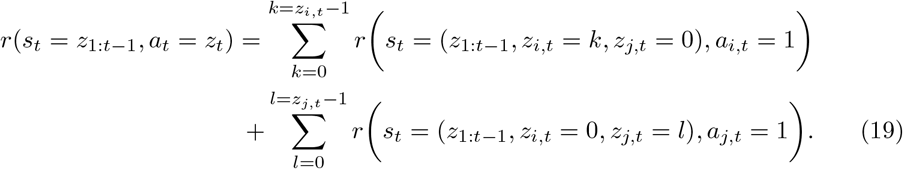

### Modeling the cost of sampling

A trade-off between the information gained from sampling (i.e. the reward) and the cost of sampling naturally arises in most settings. When optimizing sampling strategies, we therefore incorporate the costs of sampling into our reward functions by considering a net reward function:

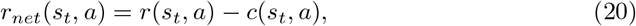

where *r*(*s*_*t*_, *a*) represents an application-specific reward function and *c*(*s*_*t*_, *a*) represents the corresponding cost of taking action *a* from state *s*_*t*_. For simplicity throughout, we assume a fixed cost per sample such that costs increase linearly with sampling effort at each time point: *c*(*s*_*t*_, *a*) ∝ *z*_*t*_.

To help ensure costs and rewards are measured on a comparable scale, we rescale both the costs and rewards relative to their maximum possible values among all policies being considered. As a result, a reward *r* = 1 implies the maximal possible amount of information has been gained. Likewise, a cost *c* = 1 implies that no more expensive strategy exists in policy space.

However, the rewards and costs are not always directly comparable on any natural scale (i.e. monetary value). Moreover, we may sometimes value the information gained from sampling more than its actual monetary cost (44). For example, we may value sampling individuals with short transmission distances more if the information we gain about transmission sources can help prevent future infections. For the transmission tree MDP, we therefore explore different *values of information v* that re-weight the relative value of the rewards relative to the costs:

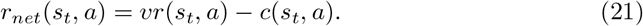

### Implementation and validation

Python code implementing all three MDPs alongside the policy evaluation and value iteration algorithms is available at:

https://github.com/davidrasm/opt-sampling-mdp. The implementation of the

MDPs was validated by comparing analytical results obtained by dynamic programming against Monte Carlo simulations of the MDP. For the exponential growth coalescent MDP, trees were simulated in msprime v1.3.3 (45). For the transmission tree distance MDP, trees were simulated in Python with the *ttd mdp*.*py* script. For the structured coalescent MDP, trees were simulated in SLiM v4.3 (46) using the script *WF assymetric migration*.*slim*. In all three cases, simulated trees were stored and manipulated as tree objects in tskit v0.6 (47). Maximum likelihood estimates of model parameters were obtained from simulated trees using previously described likelihood functions for the exponential growth (1) and structured coalescent models (4; 42).

## Results

### Optimal sampling to estimate population growth rates

We first consider how the exponential growth coalescent MDP can be used to optimize sampling strategies for estimating population growth rates. Here, we use a reward function based on Fisher information ℐ to quantify the information gained about the growth rate *γ* from the sampled data. To explore how the rewards depend on sampling times and sizes, we first consider sampling at single time points (Fig 5A). Overall, the amount of information gained about the growth rate decreases exponentially as we sample closer to the present, such that sampling more in the distant past generally provides substantially more information than sampling equivalent numbers of individuals near the present. The expected rewards computed analytically under the MDP closely track the rewards observed in Monte Carlo simulations.

**Fig 5.**
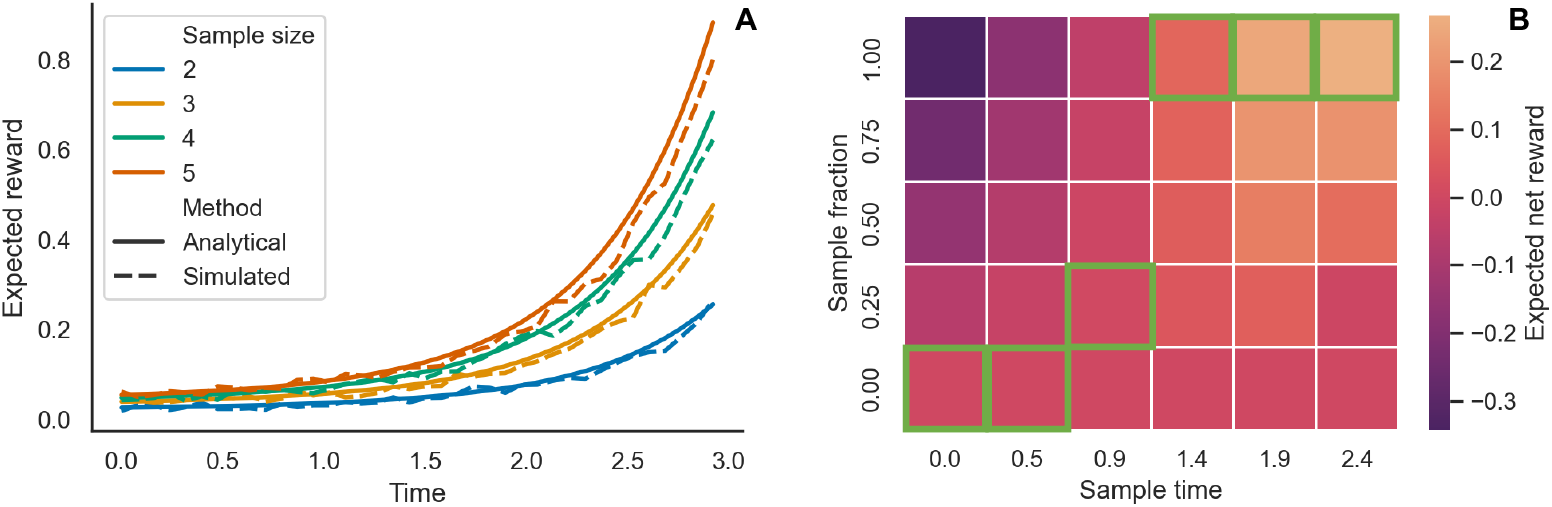
Expected rewards under the exponential growth coalescent MDP. Time is measured going backwards from the present at *τ* = 0. **A**: The expected reward or Fisher information gained from sampling different numbers of individuals at single times computed analytically (solid lines) versus the rewards returned from Monte Carlo simulations (dashed lines). **B**: The optimal sampling strategy obtained through value iteration. The optimal sampling action at each time is bordered in green. The expected net reward of each possible action is computed conditional on making the optimal action at all other times points. In all cases, *N* (0) = 50 and the growth rate *γ* = 1.0 such that time is measured in units of 1*/γ* going backwards from the present.

**Fig 6.**
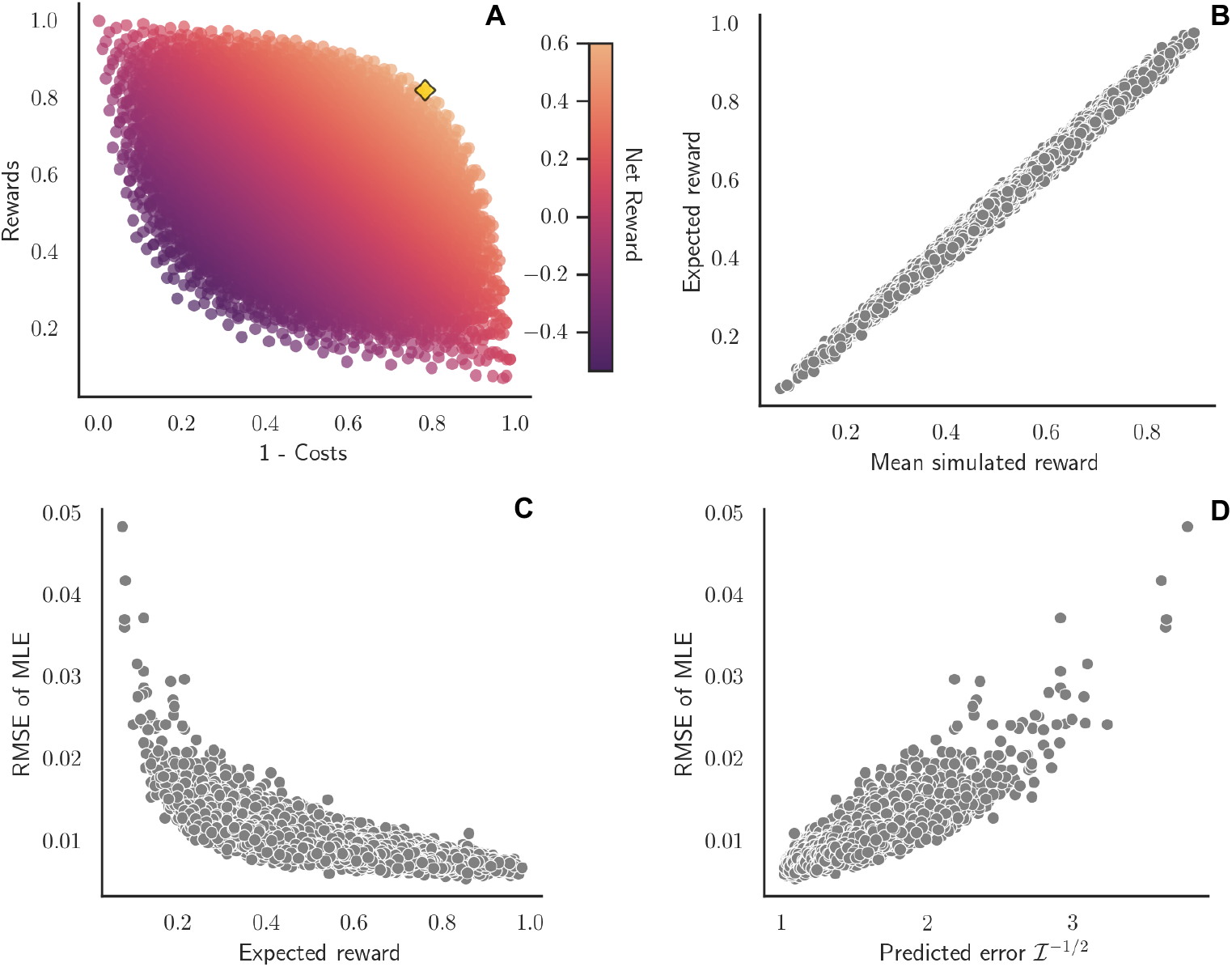
Performance of the exponential growth coalescent MDP at computing informational rewards. **A**: The rewards (Fisher information) versus costs of sampling across all 15,625 policies considered. Colors indicate the net reward returned for each policy and the optimal policy found by both the value iteration and a brute force search is marked by the gold diamond. **B**: Mean reward of each policy from Monte Carlo simulations versus expected reward computed analytically under the MDP. **C:** Expected Fisher information rewards versus root mean squared error (RMSE) of maximum likelihood estimates (MLEs) of the growth rate. **D:** Error in MLEs predicted based on expected Fisher information versus the observed RMSE.

Next, we use the value iteration algorithm to find the optimal sampling strategy when sampling sequentially through time. More specifically, we consider the problem of choosing the optimal sampling fraction from 𝒜 = {0, 0.25, 0.5, 0.75, 1.0} at six sampling time points uniformly spaced between the present and the time *τ*_*N*=1_ = log(1*/N* (0))*/r* at which *N* = 1. The optimal action in terms of sampling fractions goes from 1.0 at the earliest sampling time furthest in the past to 0.0 at present. Intuitively, this strategy is optimal because sampling earlier results in coalescent events occurring deeper in the past, which provide more Fisher information about the growth rate, while the cost of sampling higher fractions also increases as the population size grows towards present.

To ensure the optimal strategy we identified is indeed globally optimal, we performed a brute force search over all 5^6^ = 15,625 possible strategies in the policy space. For each policy, we evaluate the expected value analytically using policy evaluation and then compared this expected value against the mean reward returned from 100 Monte Carlo simulations of the MDP. Computing the reward returned by each policy against its corresponding cost reveals an apparent optimality front beyond which the informational rewards cannot be increased without paying higher costs (Fig 6A). Nevertheless, there are a large number of policies that yield high Fisher information but with relatively low cost. These policies with high net reward all concentrate sampling deep in the past. The optimal policy found by this brute force search is indeed the same as the optimal policy identified by value iteration. Moreover, rewards computed analytically under the MDP are highly correlated with the the mean reward returned by Monte Carlo simulation of each policy (Pearson correlation *ρ* = 0.996; Fig 6B).

For each Monte Carlo simulation we also obtained a maximum likelihood estimate (MLE) of the growth rate *γ* under the coalescent model. As expected, the root mean squared error (RMSE) in the MLE declines asymptotically with the expected Fisher information (Spearman rank correlation *ρ* = 0.829; Fig 6C). Because the sampling distribution of the MLEs should asymptotically converge to a normal distribution with a standard error given by the inverse of the square root of the Fisher information ℐ, we can use this relationship to predict the error in the MLEs 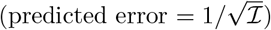. As expected, the error predicted based on Fisher information rewards accurately predict the RMSE in the MLEs (Pearson correlation *ρ* = 0.871; Fig 6D).

We further explored optimal sampling strategies under a broader range of scenarios (Table 1). The optimal strategy is invariant to the population growth rate: low (*γ* = 0.01), medium (*γ* = 0.1) and high (*γ* = 0.2) growth rates all result in the same optimal strategy. The optimal strategy changes slightly with the final population size *N* (0). Larger population sizes shift the optimal strategy to sampling even less at time points midway between the earliest and latest sampling time points. Lastly we consider scenarios where the cost of sampling decreases linearly through time, such that sampling at the earliest time is 2 to 10-fold higher than sampling at present. Such a scenario may occur, for example, if resources are limited at the beginning of an epidemic. Scenarios where the cost of sampling is 5-fold or 10-fold higher at the earliest times shift the optimal strategy slightly away from sampling at intermediate times towards sampling small fractions at present. However, the optimal strategy always remains to sample as much as possible at the earliest time points when the population is still small. Sampling towards present adds little additional value, even if the cost of sampling substantially decreases over time.

**Table 1.**
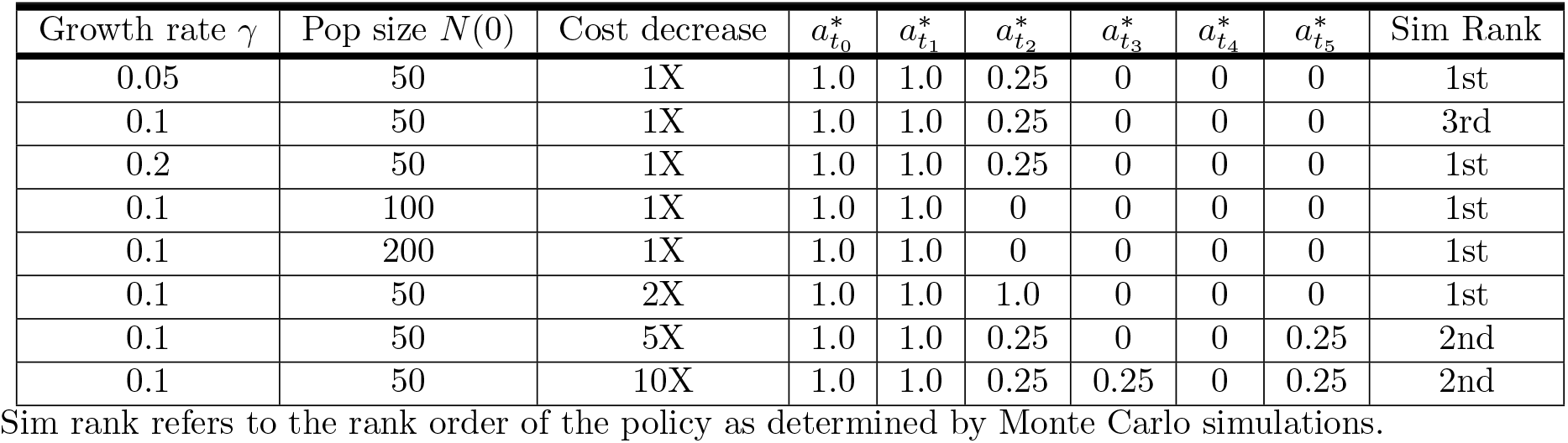
Optimal sampling strategies for estimating population growth rates. Sim rank refers to the rank order of the policy as determined by Monte Carlo simulations.

| Growth rate $\gamma$ | Pop size $N(0)$ | Cost decrease | $a_{t_0}^*$ | $a_{t_1}^*$ | $a_{t_2}^*$ | $a_{t_3}^*$ | $a_{t_4}^*$ | $a_{t_5}^*$ | Sim Rank |
| --- | --- | --- | --- | --- | --- | --- | --- | --- | --- |
| 0.05 | 50 | 1X | 1.0 | 1.0 | 0.25 | 0 | 0 | 0 | 1st |
| 0.1 | 50 | 1X | 1.0 | 1.0 | 0.25 | 0 | 0 | 0 | 3rd |
| 0.2 | 50 | 1X | 1.0 | 1.0 | 0.25 | 0 | 0 | 0 | 1st |
| 0.1 | 100 | 1X | 1.0 | 1.0 | 0 | 0 | 0 | 0 | 1st |
| 0.1 | 200 | 1X | 1.0 | 1.0 | 0 | 0 | 0 | 0 | 1st |
| 0.1 | 50 | 2X | 1.0 | 1.0 | 1.0 | 0 | 0 | 0 | 1st |
| 0.1 | 50 | 5X | 1.0 | 1.0 | 0.25 | 0 | 0 | 0.25 | 2nd |
| 0.1 | 50 | 10X | 1.0 | 1.0 | 0.25 | 0.25 | 0 | 0.25 | 2nd |

### Optimal sampling to minimize transmission distances

We next consider how to optimize sampling in order to minimize the distance between sampled individuals using the transmission tree distance MDP. To this end, we consider two different reward functions to quantify the distance between sampled individuals. Rewards based on *inverse distances* assume that the value of a sample is inversely proportional to its distance to its nearest sampled neighbor. Rewards based on *direct pairs* assume that a reward is only received for sampling direct transmission pairs.

We first consider the rewards versus costs of sampling different fractions of individuals uniformly through time in a constant sized population. Assuming the costs grow linearly with the sample fraction, the rewards grow more slowly than the costs under both reward functions (Fig 7A-B). This is especially clear when rewards are computed based on direct transmission pairs, as the expected number of direct pairs grows with the square of the sample fraction (*f* ^2^) and thus rewards grow quadratically with sampling effort.

**Fig 7.**
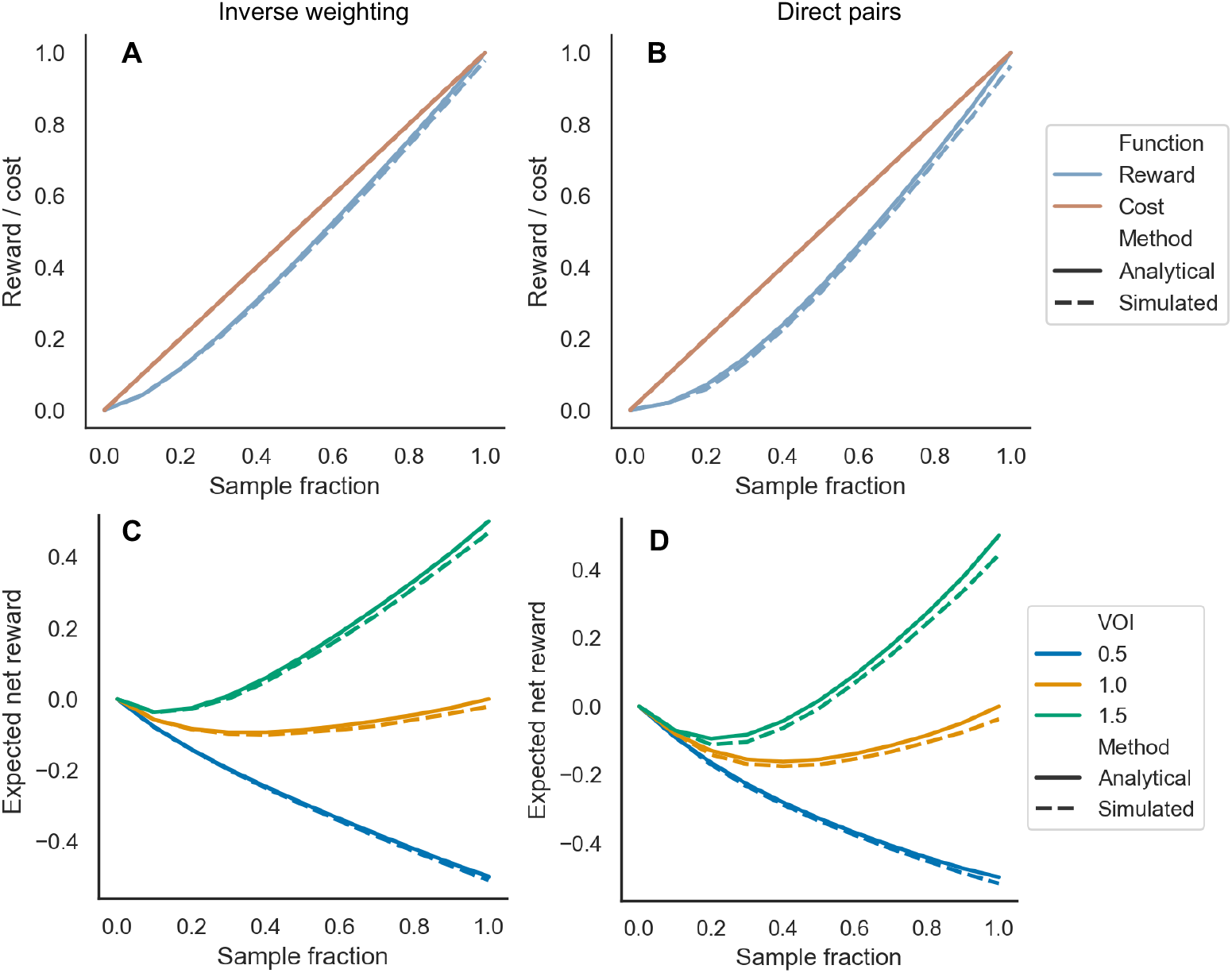
Expected rewards versus costs of sampling uniform fractions of individuals in the transmission tree MDP. In A and C, rewards are computed assuming the value of a sampled individual is inversely proportional to its distance to its nearest sampled neighbor. In B and D, rewards are computed based on the number of direct transmission pairs sampled. Rewards are computed both analytically under the MDP (solid lines) and by taking the mean across 100 Monte Carlo simulations (dashed lines). **A**: Rewards versus costs of different sampling fractions with rewards based on inverse weighting. **B**: Rewards versus costs of different sampling fractions with rewards based on direct pairs. **C**: Expected net rewards with rewards based on inverse weighting assuming different values of information (VOI), which weights the relative value of rewards versus the costs. **D**: Expected net rewards with rewards based on direct pairs.

**Fig 8.**
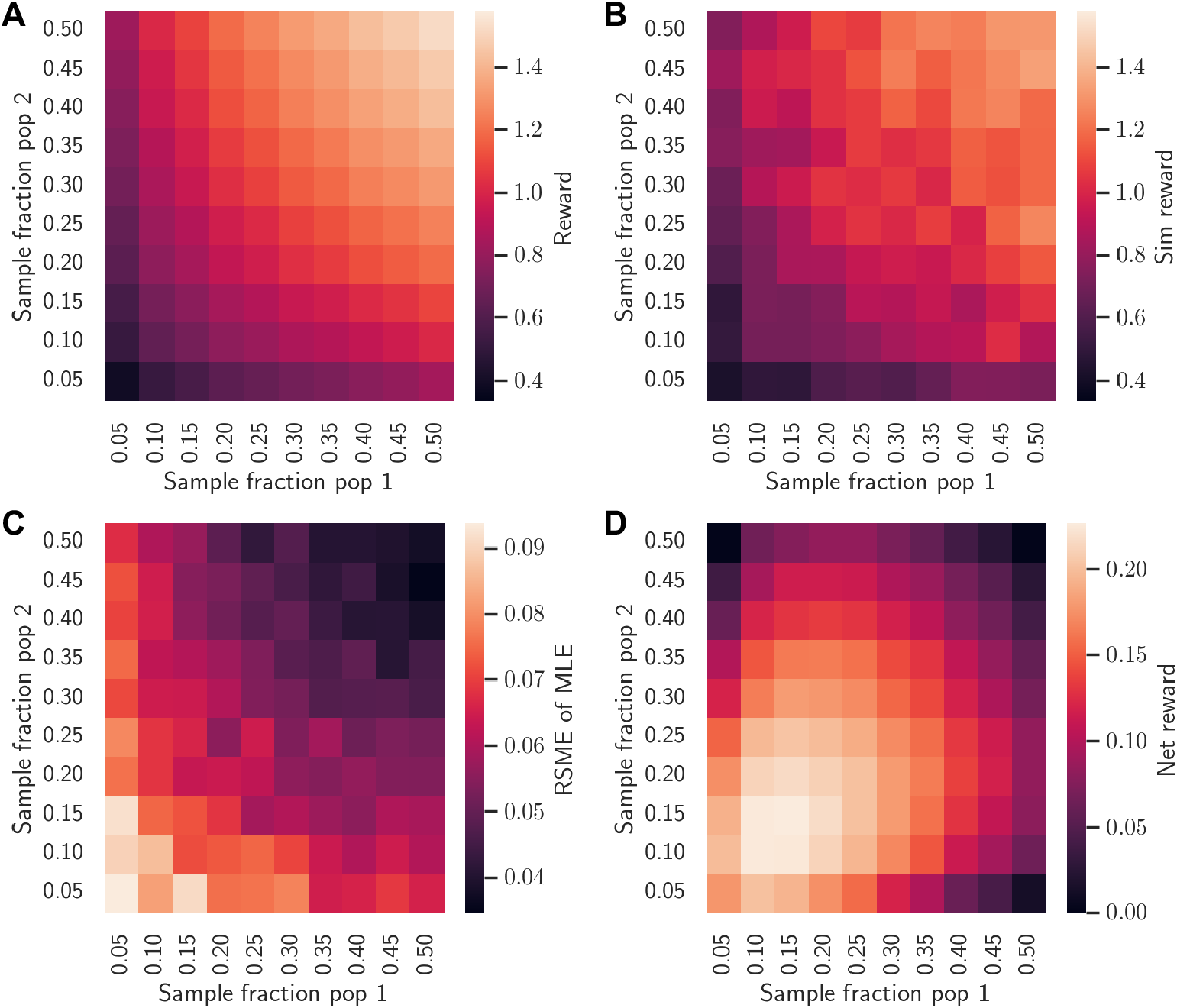
Expected rewards under the structured coalescent MDP. **A:** The expected reward of sampling varying fractions of individuals in each population at a single time point computed analytically under the MDP. **B:** The mean rewards observed from Monte Carlo simulations of the structured coalescent. In both A and B, the number of migration events is square root transformed. **C:** The average root mean squared error (RMSE) of maximum likelihood estimates (MLEs) of the migration rates from 100 simulations at each pair of sampling fractions. The reported RMSE is averaged over migration rates estimated in both directions (i.e. *RMSE* = (*RMSE*(*m*_12_)+ *RMSE*(*m*_21_))*/*2). **D:** The net reward of sampling when considering the costs of sampling. In all simulations, *N* = 50 in each population and migration is symmetric between populations (*κ* = 1.0) with the migration rates *m*_12_ = *m*_21_ = 0.02 per generation.

Assuming that the rewards are directly comparable to costs (i.e. the value of information *v* = 1.0), the net reward function is concave, with intermediate sample fractions providing the lowest net reward for both forms of the reward function (Fig 7C-D). In this case, the expected net reward of sampling either no one or everyone is very close to zero, such that both extreme sampling strategies can be considered optimal. However, when the value of information is low (*v* = 0.5), sampling no one is optimal. Conversely, if the value of information is higher (*v* = 1.5), sampling everyone becomes the optimal strategy. The optimal sampling fraction therefore strongly depends on the assumed value of information, but sampling low to intermediate fractions of individuals is generally the least optimal strategy. This is true regardless of the assumed population size because costs are computed relative to the total cost of sampling everyone.

We further explore optimal sampling strategies when sampling fractions are allowed to vary through time using value iteration. To reduce the size of action/policy space, we consider deciding among 5 sampling fractions 𝒜 = {0, 0.25, 0.5, 0.7, 1.0} at 5 consecutive generations. Even when allowing for sampling fractions to vary through time, the optimal strategy generally remains to sample either no one or everyone depending on the assumed value of information (Table 2). An exception to this general trend occurs at the earliest sampling time (deepest in the past) when the optimal action can deviate from the optimal action at other times. This is most likely due to a boundary effect created by assuming all individuals share a single non-sampled ancestor one generation before the earliest sample time. For instance, at low VOI *v* = 0.5, it may be worth sampling some individuals at the earliest time due to the fact that all individuals will necessarily share a common ancestor at the previous generation and thus have low transmission distances.

**Table 2.** Optimal sampling strategies for minimizing transmission distances. Sim rank refers to the rank order of the policy as determined by Monte Carlo simulations.

| Pop dynamics | Reward | VOI | action $t_0$ | action $t_1$ | action $t_2$ | action $t_3$ | action $t_4$ | Sim Rank |
| --- | --- | --- | --- | --- | --- | --- | --- | --- |
| Constant | Inverse | 0.5 | 0.25 | 0 | 0 | 0 | 0 | 1st |
| Constant | Inverse | 0.75 | 1 | 1 | 1 | 1 | 1 | 4th |
| Constant | Inverse | 1.5 | 1 | 1 | 1 | 1 | 1 | 1st |
| Constant | Direct | 0.5 | 0.25 | 0 | 0 | 0 | 0 | 1st |
| Constant | Direct | 1.0 | 0.5 | 1 | 1 | 1 | 1 | 5th |
| Constant | Direct | 1.5 | 0.75 | 1 | 1 | 1 | 1 | 3rd |
| Doubling | Inverse | 0.5 | 1 | 0.5 | 0 | 0 | 0 | 1st |
| Doubling | Inverse | 1.0 | 1 | 1 | 1 | 1 | 0.25 | 1st |
| Doubling | Inverse | 1.5 | 1 | 1 | 1 | 1 | 1 | 1st |
| Doubling | Direct | 0.5 | 1 | 0.5 | 0 | 0 | 0 | 1st |
| Doubling | Direct | 1.0 | 1 | 1 | 1 | 1 | 0.25 | 1st |
| Doubling | Direct | 1.5 | 1 | 1 | 1 | 1 | 1 | 1st |

We further considered optimal sampling strategies when population sizes doubled between each sampling time point, starting from a population size *N* = 1 at the first sampling time deepest in the past. Again, the optimal strategy is generally to sample no one at low VOI and sample nearly everyone at higher VOIs. The exception to this general trend occurs at the earliest sampling time points, when the population size is very small, where it may still be optimal to sample even at low VOI. Similarly, at the latest sampling time points, when the population size is large, it may be optimal to sample fewer individuals even when the VOI is high.

To ensure the optimal strategies identified by value iteration are indeed optimal, we performed a brute force search over all 5^5^ = 3,125 possible strategies under each scenario considered above using the mean reward returned by 100 Monte Carlo simulations to approximate the expected value of each policy. The optimal policy identified by a brute force search is in most cases identical to the one identified by value iteration. In cases where the optimal strategy differs there are generally multiple strategies with nearly identical expected values. Across all policies, the rank order of polices computed analytically under the MDP is strongly correlated with the rank order according to the Monte Carlo simulations (Spearman rank correlation *ρ* = 0.986).

### Optimal sampling to estimate migration rates

Finally, we consider how the structured coalescent MDP can be used to optimize sampling strategies for estimating migration or transmission rates between populations. To quantify how much information sampling provides about the migration rates, our reward function counts the expected number of coalescent events in the phylogeny of sampled individuals that occur between as opposed to within populations. As before, we first consider the simple case of sampling at a single time point, varying the fraction of individuals sampled in each population. The reward increases linearly as the sampling fraction increases in both populations and thus more coalescent events occur between instead of within populations (Fig 8A). The rewards computed analytically under the MDP closely match the mean rewards returned from Monte Carlo simulations of the structured coalescent (Fig 8B).

We also explored how the root mean squared error (RMSE) in the maximum likelihood estimates (MLEs) of the migration rates vary with the sampling fraction in both populations. Averaged across 100 Monte Carlo simulations at each pair of sampling fractions, the RMSE is generally very small unless sampling fractions are low (*f <* 0.15) in both populations (Fig 8C). We further noticed that the RMSE tends to scale with the square root of the the number of coalescent events between populations. We therefore square root transform the number of events in the reward function (see (17)). With this reward function, the optimal strategy is to sample relatively low fractions of individuals symmetrically from both populations (Fig 8D).

We further consider optimal sampling strategies when sampling fractions are allowed to vary between populations as well as across time using value iteration (Table 3). We consider deciding among *n* = 11 sampling fractions in each population, such that there are now *n*^2^ = 121 possible actions at each time. We therefore only consider sampling at three time points to limit policy space to a feasible size.

**Table 3.** Optimal sampling strategies for estimating migration rates. Sim rank refers to the rank order of the policy as determined by Monte 0043arlo simulations.

| Migration rate $m$ | Ratio $\kappa$ | $a_{p_1,t_0}^*$ | $a_{p_2,t_0}^*$ | $a_{p_1,t_1}^*$ | $a_{p_2,t_1}^*$ | $a_{p_1,t_2}^*$ | $a_{p_2,t_2}^*$ | Sim Rank |
| --- | --- | --- | --- | --- | --- | --- | --- | --- |
| 0.02 | 1.0 | 0.18 | 0.18 | 0.16 | 0.16 | 0.16 | 0.16 | 1st |
| 0.1 | 1.0 | 0.2 | 0.2 | 0.18 | 0.18 | 0.16 | 0.16 | 1st |
| 0.2 | 1.0 | 0.2 | 0.2 | 0.18 | 0.18 | 0.16 | 0.16 | 3rd |
| 0.02 | 2.0 | 0.16 | 0.2 | 0.14 | 0.18 | 0.14 | 0.18 | 1st |
| 0.1 | 2.0 | 0.16 | 0.2 | 0.14 | 0.2 | 0.14 | 0.2 | 1st |
| 0.2 | 2.0 | 0.16 | 0.2 | 0.16 | 0.16 | 0.0 | 0.2 | 4th |
| 0.02 | 10.0 | 0.1 | 0.2 | 0.12 | 0.2 | 0.1 | 0.18 | 1st |
| 0.1 | 10.0 | 0.1 | 0.2 | 0.12 | 0.2 | 0.1 | 0.2 | 1st |
| 0.2 | 10.0 | 0.1 | 0.2 | 0.12 | 0.2 | 0.08 | 0.2 | 8th |

Sim rank refers to the rank order of the policy as determined by Monte Carlo simulations.

With low and symmetric migration rates (*m* = 0.02), the optimal policy remains to sample relatively low fractions (f=0.16 - 0.18) in both populations through time. Slightly higher sampling fractions are optimal at earlier time points deeper in the past, presumably because this increases the chances of lineages sampled at later time points coalescing at events between populations. With higher but still symmetric migration rates (*m* = 0.1 and *m* = 0.2), the optimal strategy shifts slightly towards sampling higher fractions of individuals in both populations, reflecting the fact that there will on average be more migration occurring and thus more reward to be gained at higher migration rates.

We also consider optimal sampling strategies with asymmetric migration between populations, which we quantify as the ratio of the migration rates *κ* = *m*_1,2_*/m*_21_. Higher *κ* values imply migration is predominantly occurring from population 1 into 2, such that population 1 acts as source and population 2 as a sink. With moderately asymmetric migration (*κ* = 2), the optimal strategy shifts to preferentially sampling more in the sink than in the source population, especially at later time points. This strategy may be optimal because individuals sampled in the sink population are likely to have a recent ancestor and thus coalesce with lineages in the source population, whereas individuals sampled in the source population will likely coalesce with other lineages sampled in the source population, providing no additional information about migration. With highly asymmetric migration (*κ* = 10), preferentially sampling in the sink population becomes even more favored.

To ensure the sampling strategies identified by value iteration are globally optimal, we performed a brute force search over a subset of 729 out of a total of 121^3^ ≈ 1.7 million possible strategies under each scenario considered above. In almost all cases, the optimal policy identified by the brute force search is closest to the policy identified by value iteration. Ranking policies based on their total expected value or net reward, the rank order determined by policy evaluation is strongly correlated with the rank order based on Monte Carlo simulations (Spearman rank correlation *ρ* = 0.72). As expected, the the total predicted rewards are strongly negatively correlated with the RMSE in MLEs of the migration rates (Pearson correlation *ρ* = −0.61), suggesting that our reward function does indeed provide a proxy for the information gained about migration rates from simulated phylogenies.

## Discussion

By framing sampling in population genomics as a sequential decision making process, we were able to apply the well-developed machinery of MDPs to explore optimal genomic sampling strategies. The resulting MDP framework overcomes one of the main challenges in designing genomic sampling strategies—MDPs allow us to predict the expected reward or value of genomic sampling in terms of what information will be gained about estimated variables. Because the information gained from sampling will generally depend on how sampled individuals are related, we combined MDPs with coalescent models that probabilistically consider how sampled individuals might be related in a phylogeny or transmission trees to compute expected rewards. Once we can compute the expected value of sampling, the problem of designing optimal strategies reduces to finding actions or policies that maximize long-term expected rewards, which can be done very efficiently using dynamic programming.

Our MDP models provide several insights into how genomic sampling can be optimized for standard inference problems in population genomics and genomic epidemiology.

Moreover, while the models we considered are relatively simple, many of these insights appear to generalize beyond the specific details and parameters of the models. First, the exponential growth MDP shows that optimal sampling strategies for estimating population growth rates tend to concentrate sampling as early as possible, regardless of the growth rate or final size of the population. This remains true even if the costs of sampling decrease significantly over time. Samples collected early during the early stages of an epidemic are therefore likely to be most valuable for estimating other epidemiological parameters related to population growth such as *R*_0_ or the relative fitness (i.e. selection coefficient) of emerging variants (14).

Second, the transmission tree MDP reveals that sampling intermediate fractions of infected individuals can be least optimal when the goal is to minimize the transmission distance or number of intervening infections between sampled individuals. This main result appears to hold regardless of the exact assumptions about population dynamics or how transmission distances are weighted in the reward function. While this result may seem intuitive in hindsight, it highlights a potential “sunk cost fallacy” in genomic epidemiology. It may be tempting to sample more if we have already invested heavily in sampling, but increased sampling may ultimately lead diminishing returns, especially if we are starting from low sampling fractions, which is often the case in genomic epidemiology. Instead, it may be better to sample no one if the value of information (VOI) is low, or to sample everyone if the VOI is high. If the VOI fluctuates through time, an on-off or “bang-bang” strategy that switches between sampling everyone and no one may be the most pragmatic solution.

Third, the structured coalescent MDP provides insight into how genomic sampling can be optimized to infer the migration rates. The ability to infer movement is often one of the most useful applications of genomic data, as such phylogeographic approaches can shed light on the ancestry of a population or, for an infectious pathogen, possible sources of transmission such as the fraction of infections attributable to given a subpopulation (48). Reassuringly, relatively low sampling fractions across populations are often optimal even if migration is highly asymmetric. These results support other recent work demonstrating that accurate estimates of migration rates can be obtained under structured coalescent models, even when sample sizes are small and biased between populations (24).

While our simple MDPs provide insights into common sampling problems, applying our MDP framework to more complex scenarios will undoubtedly come with challenges. One major challenge lies in designing reward functions that can meaningfully capture and quantify the information gained from sampling while remaining computationally feasible to compute. When the objective is to minimize error or uncertainty about estimated parameters, rewards based on metrics such as Fisher information are a sensible and theoretically well-motivated choice, as Fisher information can be used to predict the accuracy and precision estimates prior to sampling (35; 36). In other cases, designing reward functions may be much more challenging. For instance, we only realized after much trial and error that a relatively simple reward based on the number of coalescent events occurring between populations could be used as a good proxy for information gained about migration rates.

In the long-term, MDPs will likely need to be combined with reinforcement learning (RL) methods to realize their full potential in optimizing genomic sampling. In RL, a computational agent representing a decision maker learns how to act to maximize long-term rewards by interacting with its environment (32). While MDPs provide the theoretical basis for much of RL, modern RL algorithms overcome many of the challenges inherent to learning optimal policies from MDPs in complex environments. Perhaps most importantly, RL algorithms do not require perfect knowledge of the underlying dynamics of the system (i.e. the Markov process) since they learn to act optimally based purely on experience. In settings where there may be no simple mapping between actions and rewards, RL can be further combined with Q-learning methods to learn how to predict q-values or the expected reward received from taking a given action from a given state (49; 50). These Q-learning approaches could then be used to optimize genomic sampling by training the agent to predict what new information will be gained from sampling and then acting accordingly.

Going forward, the simple MDP models presented here can be extended to address many of the challenges confronting genomic sampling design in real-world applications. Potential applications include population dynamics beyond exponential growth, which may require adaptive sampling through time (51). In genomic epidemiology, the transmission tree MDP could be extended to include unobserved or asymptomatic infections that would require optimizing testing strategies alongside optimizing genomic sequencing strategies. In phylogeography, our two-population structured coalescent model could be extended to consider the movement of individuals through many subpopulations or across complex landscapes. In each of these scenarios, MDPs can provide analytical insights in simple cases, which can then be built upon by applying RL methods to confront more complex scenarios.

## Supporting information

**S1 Appendix. Optimization using dynamic programming**. Provides pseudocode for the policy evaluation algorithm used to evaluate the value of a given policy and the value iteration algorithm for optimizing policies using dynamic programming.

**S2 Appendix. Exponential growth coalescent MDP**. Describes the details of how the coalescent probability density is computed under the exponential growth coalescent model to compute expected rewards.

**S3 Appendix. Transmission tree distance MDP**. Describes the details of how the probability of a sampled individual having a nearest sampled neighbor at some distance in a transmission tree is computed.

**S4 Appendix. Structured coalescent MDP**. Describes the details of how the coalescent probability density is computed to find expected rewards under the structured coalescent model.

## Acknowledgments

This work was supported by the U.S. Center for Disease Control and Prevention award #U01CK000587-01 and the Research Capacity Fund (HATCH) project award 7003207 from the U.S. Department of Agriculture’s National Institute of Food and Agriculture.

We would also like to thank Dr. Cristina Lanzas and the members of the CDC Modeling Infectious Diseases in Healthcare Network (MinD) Whole Genome Sequencing working group for conversations that inspired this work.

## Supporting Information

## 1 Optimization using dynamic programming

To evaluate the expected value of a given policy *π*, we use an iterative policy evaluation algorithm (1). Starting from an initially arbitrary estimate of the value *V* (*s*_*t*_) for each state *s*_*t*_ ∈ 𝒮, each iteration of the algorithm updates the estimate of *V* (*s*_*t*_) using the expected value of successor states *V* (*s*_*t*+1_) obtained from previous iterations. This procedure is iterated until the algorithm converges on a stable estimate of *V*.

### Algorithm 1

Iterative Policy Evaluation

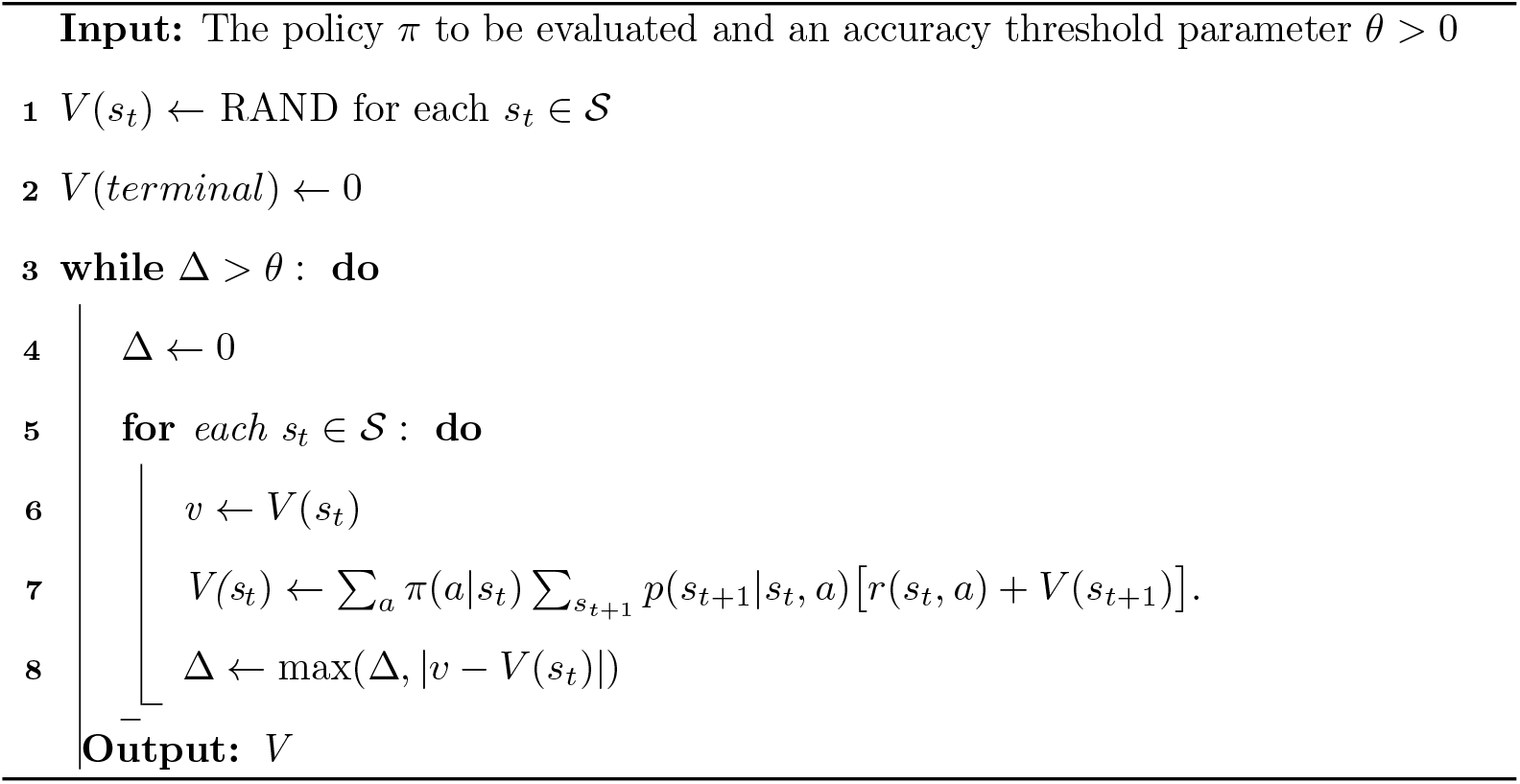

To identify the optimal policy, we use a value iteration algorithm (2; 1). This algorithm combines iteratively updating expected values with improvements to the policy. This procedure is iterated until the algorithm converges on a stable estimate of *V* where the policy can no longer be improved. Given the optimized value function *V*, we can then find the optimal action *a*^*∗*^ to take from any state *s*_*t*_: *a*^*∗*^ = arg max_*a*_[*q*_*π*_(*s*_*t*_, *a*)]. Identifying *a*^*∗*^ for all possible states therefore provides the optimal policy.

### Algorithm 2

Value Iteration

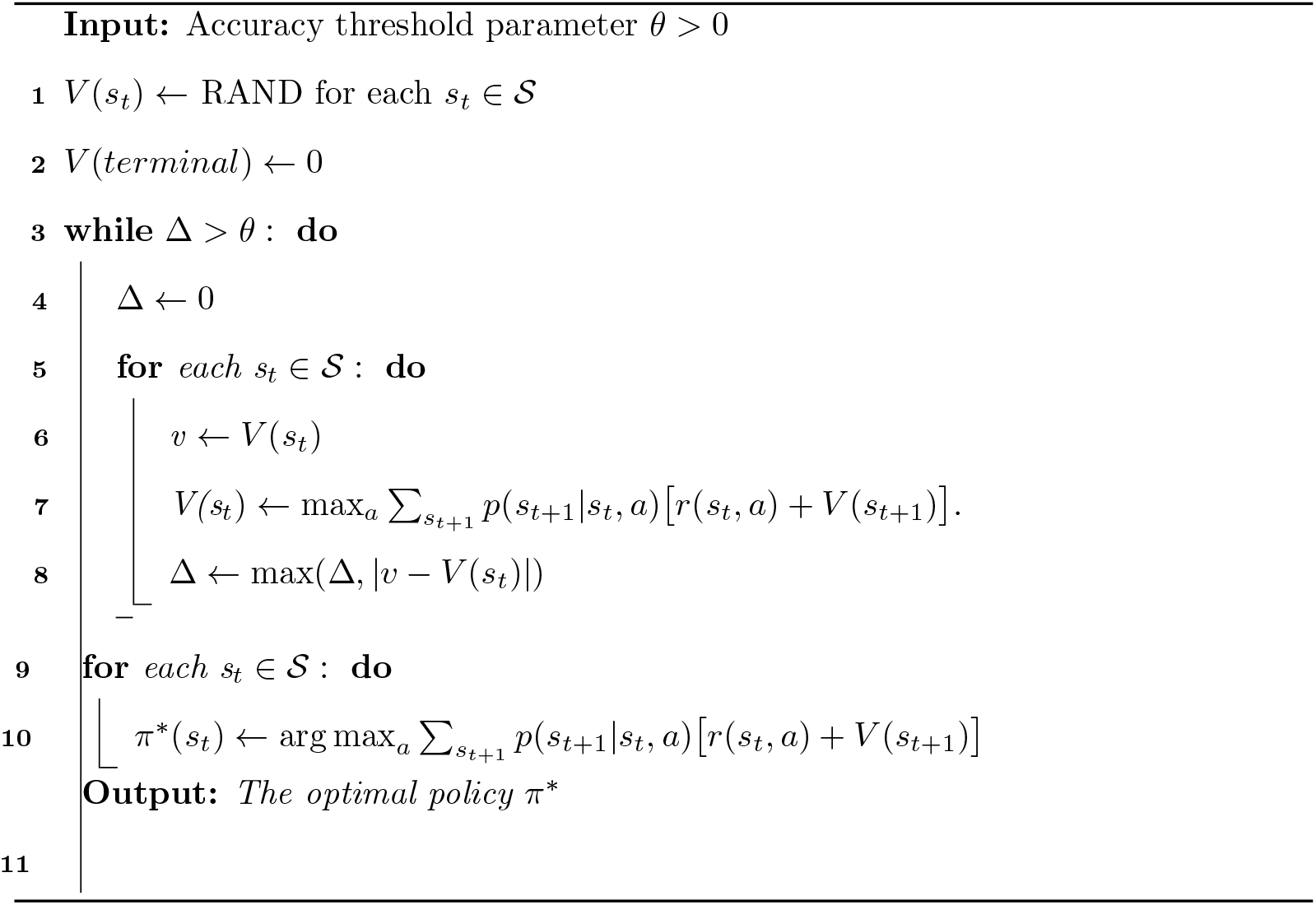

## 2 Exponential growth coalescent MDP

In order to compute the expected value of a new sample under the exponential coalescent model, we track the probability density *p*(*τ* |*z*_1:*t*_) for the time at which the newly sampled individual coalesces with the other sampled lineages, conditional on the sample configuration *z*_1:*t*_.

### 2.1 Lineages through time density

To obtain *p*(*τ* |*z*_1:*t*_), we first need to consider the lineage through time (LTT) density *q*(*l, τ*), which gives the probability that *l* other sampled lineages are present at time *τ* in the past.

In a single generation, the probability that a single pair of lineages coalesces is 1*/N* (*τ*), such that the probability a given pair does not coalesce is 1 − 1*/N* (*τ*). With *k* lineages present, there are 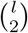 pairs of lineages which could potentially coalesce. Assuming more than one coalescent event cannot occur per generation, the transition probabilities for how the number of lineages changes between generations are:

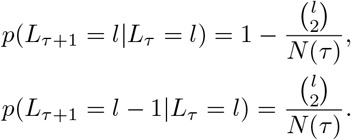

We can iteratively compute *q*(*l, τ*) backward in time given these transition probabilities, or we can approximate these probabilities in continuous-time by solving the following system of differential equations:

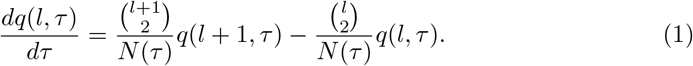

Sequential sampling through time can be accommodated by updating *q*(*l, τ*) at sampling events to reflect the addition of new lineages. If at time *τ* in the past *z* lineages are sampled, we update *q*(*l, τ*) as:

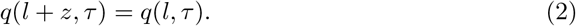

Numerically solving *q*(*l, τ*) through time shows that the computed lineage through time probabilities match the trajectories of lineages through time in Monte Carlo simulations of the coalescent process (Fig 1A).

### 2.2 Coalescent time probability density

As originally shown in (3), the probability that a pair of sampled lineages coalesce at time *τ* in the past in an exponentially growing population is:

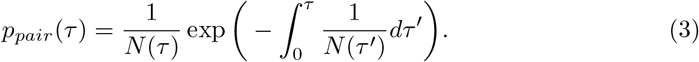

Modifying this density to take into account any possible number of sampled lineages *l* through time, the probability density *p*(*τ* |*z*_1:*t*_) for the time at which a sampled lineage coalesces with the rest of the sample is:

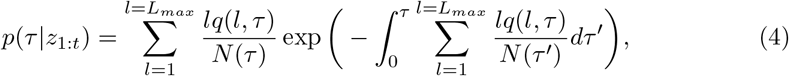

Where 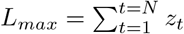, the maximum possible number of samples in the tree.

Fig 1B shows the probability density for the time at which a newly sampled lineage coalesces with the rest of the sample for the case where four other lineages are sampled at present. The analytical density given by Eq (4) closely follows the observed distribution of coalescent times from Monte Carlo simulations.

## 3 Transmission tree distance MDP

In order to compute the expected value of a given sampling action in the transmission tree MDP, we track two auxiliary probability densities backwards through time. The first tracks the number of individuals or lineages ancestral to the sample at each generation. We refer to this as the lineages through time (LTT) density. The second tracks the transmission distance between a representative ancestor and its nearest sampled descendant. We refer to this as the ancestor-to-sample distance density.

Note that, as a shorthand below, we refer to any individual that has a sampled descendant as a sampled ancestor, regardless of whether that individual is directly sampled or not.

### 3.1 Lineages through time density

Let *q*(*l, t*) be the probability that *l* sampled ancestors are present in the transmission tree at time *t*, conditional on the number of samples taken at each generation *z*_1:*t*_. Going backwards in time, the number of sampled ancestors can change due to coalescent events where two or more sampled ancestors share the same parent in the previous generation.

Given *L*_*t*_ = *k* sampled ancestors at time *t*, the probability that there are *L*_*t*+1_ = *l* sampled ancestors one generation in the past is:

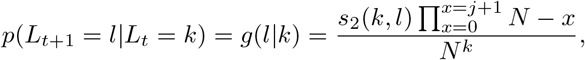

where *s*_2_(*k, l*) is the Stirling number of the second kind and *N* is the population size (4; 5). Note that these transition densities account for the possibility of multiple lineages coalescing in a single generation, which may be likely when the number of sampled ancestors is large relative to *N*.

Given the transition densities *g*(*l*|*k*), we can update *q*(*l, t*) at each generation going into the past using the following recursion:

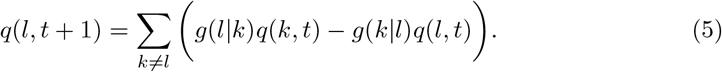

The number of sampled ancestors can also change due to sampling events at each generation. However, not all samples will necessarily increase the number of individuals ancestral to sample because a newly sampled individual may already be a sampled ancestor. In other words, the sampled individual may already have a sampled descendant in the sample. Assuming that there are *l* sampled ancestors prior to the sampling event, the probability that a randomly sampled individual is already a sampled ancestor is *p*_*anc*_ = *l/N*. Given *z*_*t*_ new samples, the probability that *k* of these individuals are newly sampled ancestors not already in the sample follows a binomial distribution:

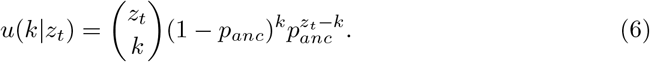

We can therefore update *q*(*l, t*) for all possible values of *l* at sampling events by considering how many of the *z*_*t*_ sampled individuals are newly sampled ancestors:

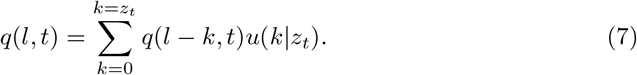

Solving the recursion for *q*(*l, t*) backwards through time shows that the lineage through time probabilities match the equivalent probabilities computed from simulated transmission trees (Fig 2A).

### 3.2 Ancestor-to-sample distance density

We also track the transmission distance between a representative sampled ancestor and its nearest directly sampled descendant. Conditional upon an individual being a sampled ancestor, let *m*(*k, t*) be the probability that the transmission distance between this individual and its nearest sampled descendant is *k*. For a newly sampled individual *m*(*k, t*) = 0. Without further sampling, the transmission distance between our representative ancestor and its sampled descendant increases by one each generation going into the past, such that *m*(*k* + 1,*t* + 1) = *m*(*k, t*) for all *k*.

However, if we are sampling sequentially though time, the distance of our representative ancestor to its nearest sampled descendant will reset to zero if it is sampled again at an earlier generation (i.e. if it is resampled). We therefore need to compute the probability that our representative ancestor is among the individuals sampled at a sampling event. This is complicated by the fact that we do not necessarily know the total number of sampled ancestors in the transmission tree before the sampling event (but only *q*(*l, t*)) nor the number of new sampled ancestors the sampling event will add to the sample (because we may resample any already sampled ancestor). Nevertheless, we can compute the overall probability that our representative ancestor is among the *z*_*t*_ newly sampled individuals by summing over all possible numbers of total ancestors *l* and newly sampled ancestors *k*:

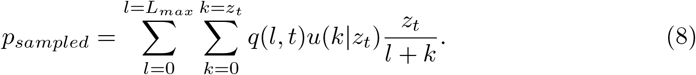

We then update *m*(*k, t*) after the sampling event by setting *m*(0, *t*) = *p*_*sampled*_ and *m*(*k, t*) = (1 − *p*_*sampled*_)*m*(*k, t*) for *k >* 0.

The distance of our representative ancestor to its nearest sampled descendant may also change if it coalesces with another lineage that has a closer sampled ancestor. To account for this possibility, we first compute that probability that our representative ancestor coalesces with any of the other *l* sampled ancestors in a given generation:

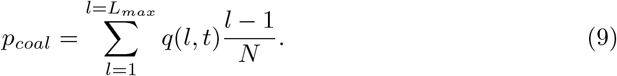

Conditional upon our representative ancestor coalescing with another sampled lineage, let *m*(*l, t*) be the distance of the other lineage to its nearest sampled descendant. Assuming all lineages in the sample are exchangeable, *m*(*l, t*) = *m*(*k, t*). The probability that the other lineage is closer to its nearest sampled descendant than our representative ancestor is therefore 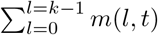. Likewise, the probability that the other lineage is further away from its nearest sampled descendant is 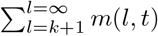. We can then combine these probabilities to update *m*(*k, t*) at each generation going into the past using the following recursion:

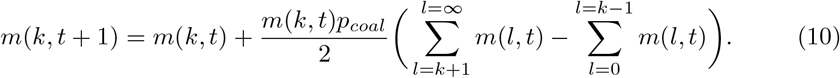

Note that we divide by two here since the probability that our representative ancestral lineage is the lineage with the lower of the two distances is 1*/*2.

Solving the recursion for *m*(*k, t*) backwards through time gives mean ancestor-to-sample distances. The resulting distance strongly agree with the mean distances computed from simulated transmission trees (Fig 2B).

### 3.3 Transmission distance probabilities

Given *q*(*l, t*) and *m*(*k, t*), we can combine these two probability densities to compute the probability *v*(*k*|*Ƶ*_1:*t*_) that a newly sampled individual had a nearest sampled neighbor in *Ƶ*_1:*t*_ at any distance *k*.

To compute *v*(*k*|*Ƶ*_1:*t*_), we first consider the probability *w*(*k*) that a sampled individual *I* has a sampled neighbor *j* at some transmission distance *d*(*i, j*) = *k*. In order for *d*(*i, j*) = *k, i* and *j* must share a common ancestor *h* at some *t*_*coal*_ where the transmission distances *d*(*i, h*) and *d*(*j, h*) sum to *k*. We can therefore think about *k* as being the sum of two random variables, *x* = *d*(*i, h*) and *y* = *d*(*j, h*)), determined by the unobserved topology of the transmission tree.

If individual *i* is sampled at time *t*_*sample*_, it must coalesce with another sampled lineage at time *t*_*coal*_ = *t*_*sample*_ + *x* in order for *x* = *d*(*i, h*). The probability that lineage *i* coalesces with another sampled lineage at *t*_*coal*_ is:

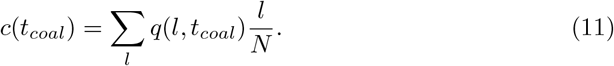

Given a coalescent event at this time, the probability that the other child lineage has a sampled descendant at transmission distance *y* = *d*(*j, h*) is provided by *m*(*y, t*_*coal*_).

To compute *w*(*k*), we can then sum the probability of all coalescent events for which *x* + *y* = *d*(*i, h*)+ *d*(*j, h*) = *k*:

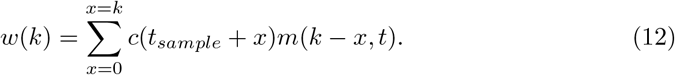

However, *w*(*k*) simply represents the probability that a sampled individual has a sampled neighbor at transmission distance *k*. To compute the probability that a sampled individual’s *nearest* sampled neighbor is at transmission distance *k*, we need to to consider the (cumulative) probability that the sampled individual does not have a sampled neighbor at a distance less than *k*, but does have a sampled neighbor at distance *k*. We therefore arrive at our desired probability density:

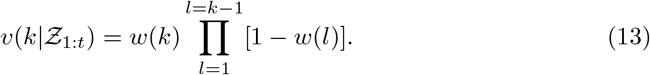

The probability density *v*(*k*|*Ƶ*_1:*t*_) strongly agrees with probabilities of a randomly sampled individual having a nearest sampled neighbor at distance *k* in simulated transmission trees (Fig 2C).

## 4 Structured coalescent MDP

In order to compute the expected value of a new sample under the structured coalescent, we track the joint probability density *p*(*τ, u, v*|*z*_1:*t*_) for the time *τ* in the past at which the sampled individual coalesces with the rest of the sample along with the state *u* of the parent lineage and state *v* of the child (sampled) lineage at the coalescent event. As with the exponential coalescent MDP, computing the coalescent density *p*(*τ, u, v*|*z*_1:*t*_) requires tracking the number of lineages through time (LTT) density backwards through time, but now for the number of lineages in each population.

### 4.1 Lineages through time density

For the structured coalescent model with two populations, let *q*(*k, l, τ*) be the probability that *k* sampled lineages are in population 1 and *l* sampled lineages are in population 2 at time *τ* in the past. To track the number of lineages through time, we distinguish between three different types of coalescent events, as each event type causes the number of lineages to change differently.

- **Type 1:** Coalescent events between lineages in the same population. These events will cause the number of lineages to decrease by one in that population.
- **Type 2:** Coalescent events between lineages in different populations. In this case, one of the two child lineages will be in the same population as the parent lineage, while the other child will be in the other population. The number of lineages will decrease by one in the non-parental population.
- **Type 3:** Unobserved coalescent events arising from one of the two child lineages not being sampled. If the sampled child lineage is in a different state from its parent, the sampled lineage moving between populations at an unobserved coalescent event. The number of lineages in the child population will go down by one and go up by one in the parental population.

Let *L*_*τ*_ = (*k, l*) denote the configuration of lineages at time *τ*, where *k* is the number of sampled lineages in population 1 and *l* is the number in population 2. Given *L*_*τ*_, we consider the probability of transitioning to *L*_*τ*+1_ at time *τ* + 1 one generation in the past. Assuming more than one event cannot occur per unit of time, the transition probability densities are:

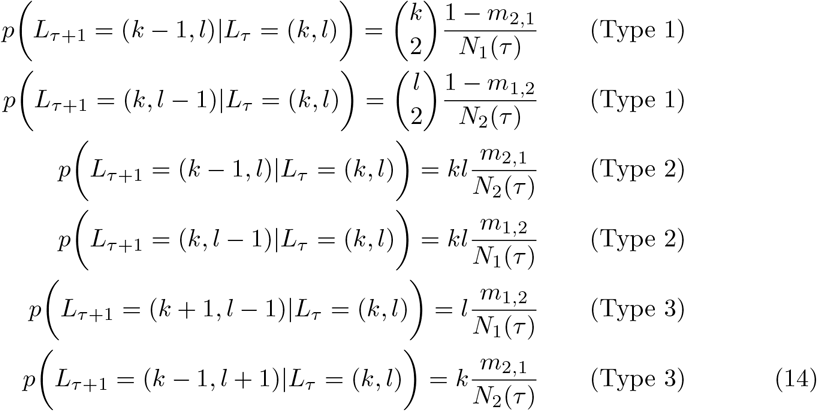

Based on these transition densities, we obtain the the following system of differential equations to track how *q*(*k, l, τ*) evolves backwards in continuous time:

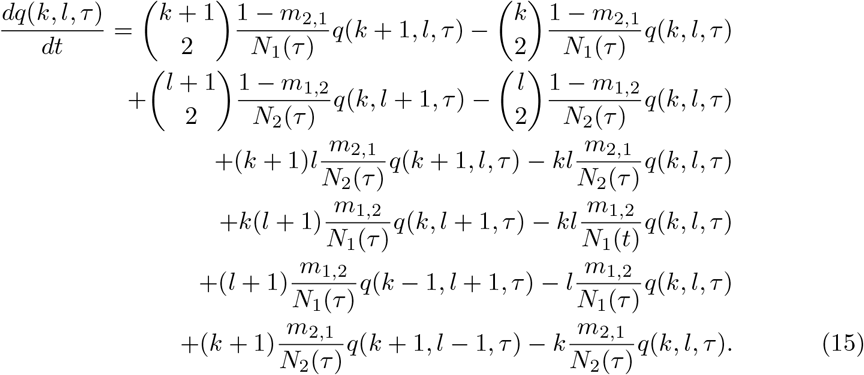

Numerically solving *q*(*k, l, τ*) through time shows that the expected trajectory of lineages in each population through time matches the trajectories in Monte Carlo simulations of the structured coalescent process (Fig 3A).

### 4.2 Structured coalescent density

The probability density *p*(*τ, u, v*|*z*_1:*t*_) can be computed iteratively at each time step in a three step process.

First, we compute the expected number of lineages in each population *A*_1_(*τ*) and *A*_2_(*τ*) at time *τ*. These expected values can be computed from *q*(*k, l, τ*) by marginalizing over all possible configurations of lineages between the two populations:

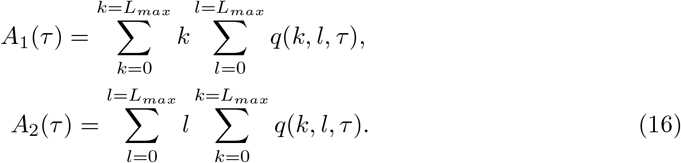

Second, we track the probabilities *s*_1_(*τ*) and *s*_2_(*τ*) that the lineage ancestral to the newly sampled individual resides in population 1 or 2, respectively. The probability that the sampled lineage resides in each population can then be tracked using the following coupled pair of differential equations:

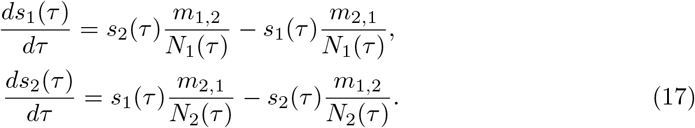

Note that here we only consider the probability of the lineages moving between states due to unobserved (Type 3) coalescent events since Type 1 and 2 events would be observed in the phylogeny and we are only interested in what population the sampled lineage resides in before it coalesces with another sampled lineage.

Third, given *A*(*τ*) and *s*(*τ*), we compute the rate *λ*_*u,v*_(*τ*) at which the sampled lineage coalesces at an event where the parent lineage is in state *u* and the child (sampled) lineage is in state *v*.

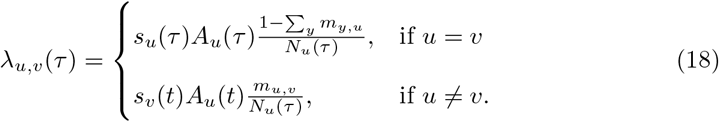

The coalescent density *p*(*τ, u, v*|*z*_1:*t*_) is then obtained by considering the probability that no coalescent event occurs prior to time *τ* along with the probability that the sampled lineage coalesces with a parent in lineage in state *u* while in state *v* at time *τ* :

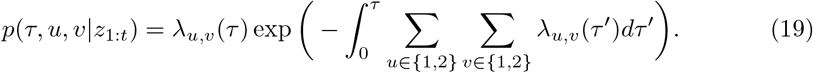

Fig 3B shows the probability density for the time at which a newly sampled lineage coalesces with a parent lineage in each population for the case where four other lineages (two in each population) are sampled at present. The density given by Eq (19) closely follows the observed distribution of coalescent times in each population observed in Monte Carlo simulations of the structured coalescent.

